# A Broad Survey and Functional Analysis of Immunoglobulin Loci Variation in Rhesus Macaques

**DOI:** 10.1101/2025.01.07.631319

**Authors:** Ayelet Peres, Amit A. Upadhyay, Vered Klein, Swati Saha, Oscar L. Rodriguez, Zachary M. Vanwinkle, Kirti Karunakaran, Amanda Metz, William Lauer, Mark C. Lin, Timothy Melton, Lukas Granholm, Pazit Polak, Samuel M Peterson, Eric J Peterson, Nagarajan Raju, Kaitlyn Shields, Steven Schultze, Thang Ton, Adam Ericsen, Stacey A. Lapp, Francois Villinger, Mats Ohlin, Christopher A. Cottrell, Rama R. Amara, Cynthia A. Derdeyn, Shane Crotty, William R. Schief, Gunilla B. Karlsson Hedestam, Melissa L. Smith, William Lees, Corey T. Watson, Gur Yaari, Steven E. Bosinger

## Abstract

Rhesus macaques (RMs) are a vital model for studying human disease and invaluable to pre-clinical vaccine research, particularly for the study of broadly neutralizing antibody responses. Such studies require robust genetic resources for antibody-encoding genes within the immunoglobulin (IG) loci. The complexity of the IG loci has historically made them challenging to characterize accurately. To address this, we developed novel experimental and computational methodologies to generate the largest collection to date of integrated antibody repertoire and long-read genomic sequencing data in 106 Indian origin RMs. We created a comprehensive resource of IG heavy and light chain variable (V), diversity (D), and joining (J) alleles, as well as leader, intronic, and recombination signal sequences (RSSs), including the curation of 1474 novel alleles, unveiling tremendous diversity, and expanding existing IG allele sets by 60%. This publicly available, continually updated resource (https://vdjbase.org/reference_book/Rhesus_Macaque) provides the foundation for advancing RM immunogenomics, vaccine discovery, and translational research.

## 1 Introduction

The extraordinary diversity of B cell receptors (BCRs) underpins the adaptive immune system’s ability to recognize a vast range of foreign antigens and produce high-affinity, neutralizing antibodies [1]. In mammals, BCRs are encoded by an extensive gene family that reside within three immunoglobulin (IG) loci: the heavy chain locus (IGH), and the light chain loci, kappa (IGK) and lambda (IGL). Diversity in the expressed BCR repertoire is generated through the recombination of V, D (IGH only), and J gene segments, combined with imprecise joining of the segments during B cell development. This diversification process is further enhanced by somatic hypermutation (SHM) after antigen exposure [2]. Despite their fundamental role in forming the BCR repertoire, the study of germline IG gene diversity is in its infancy [3]. Historically, the IG loci have been difficult to sequence and characterize using standard high-throughput genomic approaches, largely due to their genomic complexity, represented by high repeat content and segmental duplication, compounded by extensive genetic polymorphic diversity between individuals. Recent advances in genomic and expressed BCR repertoire sequencing and analysis methods have enabled large-scale characterization of the IG loci, uncovering the first hints of the remarkable allelic and haplotype diversity present at the population level [4, 5, 6, 7]. Importantly, there is a growing appreciation for the fact that the composition of an individual’s BCR repertoire is shaped at multiple levels by heritable factors within the IG loci [8, 9, 10, 5], and even single nucleotide polymorphisms resulting in coding changes can alter the ability of an IG gene to encode protective IGs [11, 12, 13, 14].

Many efforts toward cataloging IG genes and alleles in rhesus macaques (RMs) have emerged over the past decade [15, 16, 17, 18, 19, 20, 21, 22, 6, 23], providing initially valuable resources for immunogenetic analyses, and highlighting the immense diversity of the RM IG loci. Together, these studies have used genomic and expressed BCR sequencing data to characterize IG sequences. Analysis of short-read and long-read sequencing-based assemblies has revealed that RM IG loci are structured similarly to those of humans, each spanning large ĩMB regions of the genome and harboring hundreds of functional/open reading frame variable (V), diversity (D), and joining (J) gene segments [18, 19, 22]. Although these studies provided an initial impression of locus structure, they were based on assemblies from only a small number of individual RMs and of variable and undefined quality. However, initial comparisons of these haplotypes revealed the presence of large structural variants (SVs), altering gene copy number, and highlighting the potential for significant haplotype variation [19, 18]. Additional IG germlines have been identified from expressed BCR and targeted genomic sequencing [21, 17, 6, 7, 23] in additional animals, from which hundreds of allelic variants have been curated. These studies have highlighted the tremendous genetic diversity of IG present in RM, providing early findings that this diversity is similar to, and likely exceeds, that known for humans.

While the RM IG genetic studies conducted to date have been instrumental in establishing essential foundational resources for the community, they currently face several limitations: (1) they are collectively heavily biased toward IGHV sequences, with limited coverage of light chain alleles; (2) with the exception of Vazquez Bernat et al., [6] they are derived from small subsets of animals, offering limited representation of colony and species diversity; (3) they have been sourced largely from expressed BCR sequencing datasets, lacking information on locus position and annotation of elements, such as introns, leaders, and recombination signal sequences (RSSs). This last limitation, specifically, has impeded efforts to carry out comprehensive gene and allele assignments for the majority of known IG germlines; as a result, alleles identified across studies and available resources lack a unified nomenclature system. The importance of continuing to build more robust IG germline resources is motivated by the fact that RM remains a cornerstone model in biomedical and pre-clinical research [24, 25, 26]. RMs of Indian origin, in particular, are valuable for the preclinical testing of candidate HIV vaccines [27]. This is highly relevant, as many of the most promising HIV vaccine candidates aim to elicit B cell responses from a predefined set of IG genes, a strategy known as “germline-targeting” (GT) [28]. This approach focuses on the elicitation of broadly neutralizing antibody (bNAb) precusors encoded by specific germline (unmutated) IG gene segments. In humans, it has been clearly demonstrated that the IG genotype and germline BCR precursor frequencies in the pre-vaccination repertoire can significantly influence vaccination outcomes [29, 30]. This observation necessitates a clear understanding of putative RM homologs for these candidate GT vaccine targets in human. Thus, the paucity of comprehensive and well-curated IG genomic resources has the potential to constrain the utility of RMs for evaluating GT immunogens. This potential limitation is illustrated by studies in which the GT immunogen eOD-GT8 successfully elicited VRC01-class B cell precursors in humans but failed to do so in RM, likely due to the absence of a direct homolog to the human IGHV1-2 gene [29, 31]. Conversely, GT immunogens designed for IG genes with closely related homologs in both species have successfully elicited B cells using the appropriate IG gene segment in RM [32]. As additional GT immunogens advance through preclinical development [33], a comprehensive understanding of the IG loci in RM has become critical.

To better meet the growing needs of the community, we initiated the RM IG Sequencing Project, which aims to conduct population-scale genotyping of the IGH, IGK, and IGL loci in RMs from AIDS-designated colonies throughout the US. In order to overcome the technical hurdles associated with sequencing and annotation of these notoriously challenging regions, we implemented a novel approach of re-constructing IG haplotype-resolved assemblies utilizing long-read sequencing of genomic DNA and pairing this data with expressed adaptive immune receptor repertoire sequencing (AIRR-Seq) analyses. Here, we report the first release of alleles from this project, resulting from paired analyses of 106 Indian origin RMs sourced from three different colonies. Curated datasets have been made publicly available; they provide a significantly expanded resource for immunogenetic analyses of the RM, and lay the foundation for a unified nomenclature system of the IG loci in this species.

## 2 Results

### 2.1 Unprecedented Collection of IG Alleles Illuminates Surprisingly High Levels of Germline Variation

To assess the diversity of the RM IG loci, we collected PBMCs from 106 RMs in three different primate centers throughout the US and generated matched IGH, IGK, and IGL targeted long-read genomic (DNA) sequencing data and expressed (RNA) IgM, IgK, and IgL BCR repertoires (Supplementary Table 1) (Figure 1). This dataset served as the starting point for curation of an unprecedented collection of IG-encoding alleles in RM, and enabled an in-depth exploration of IG germline variability using the strategy depicted in Figure 1.

**Figure 1:**
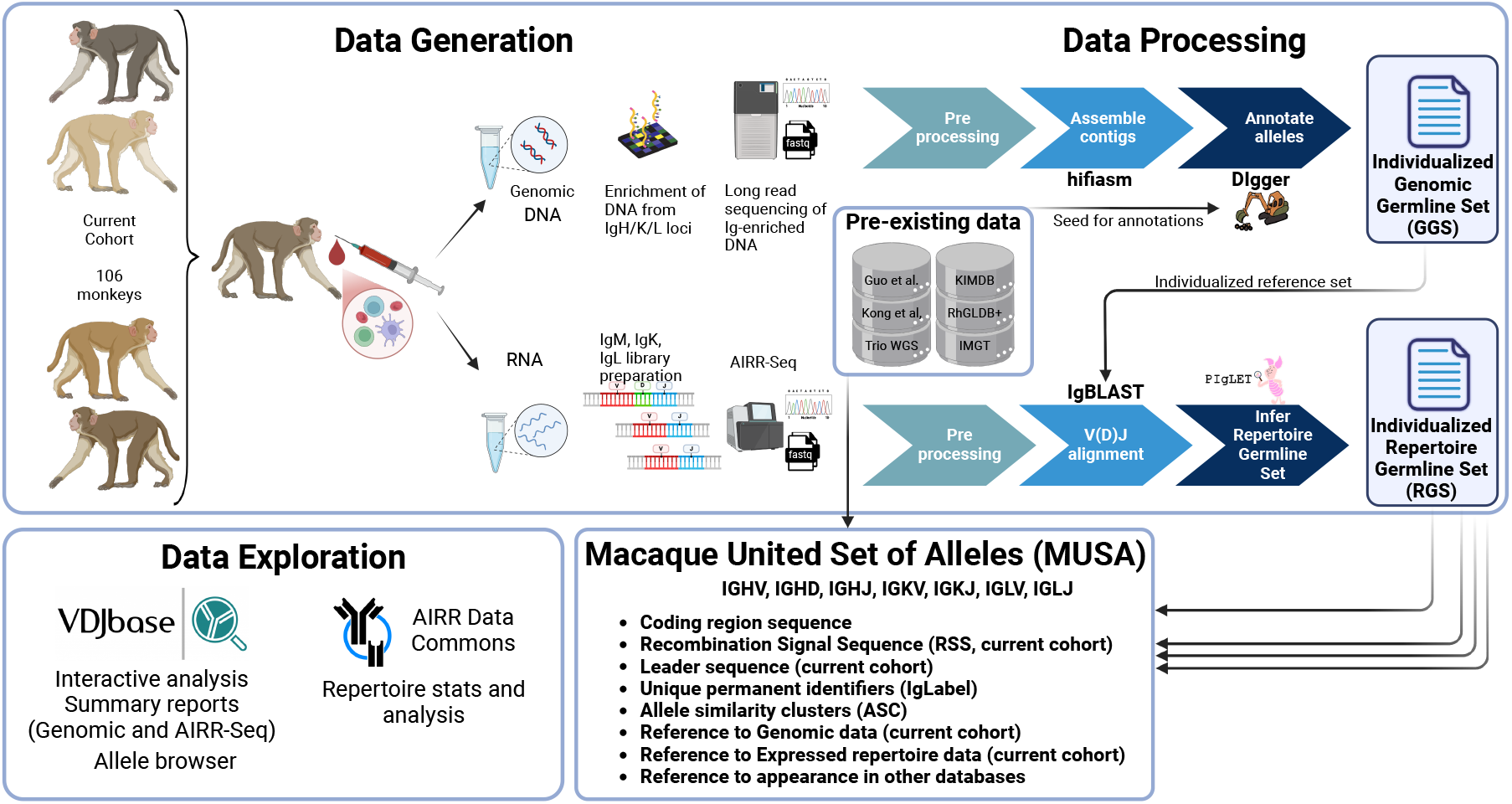
Overview of sample collection, data generation, bioinformatics and public deposition of the RM IGSeq project. PBMCs were collected from 106 RMs for RNA and DNA extraction from three primate centers across the US, and RNA and genomic DNA were extracted for each animal. Genomic DNA from IG-encoding loci was enriched and sequenced using PacBio long-read sequencing. To serve as a reference for annotating putative alleles generated by DNA genomic data we generated a provisional set of alleles (the Baseline Set) by amalgamating alleles from published sets (KIMDB[6], RHGLDB[21], IMGT [22], Kong et al. [38], and Guo et al.[23]). DIgger was used to annotate alleles, generating an individualized Genomic Germline Set (GGS) for each animal. To generate expressed B cell receptor repertoire (AIRR-Seq) data for each loci, IgM, IgK and IgL libraries were generated from RNA and sequenced using Illumina sequencing. AIRR-Seq data for each loci for each individual was mapped against the individualized GGS for each animal using IgBLAST. Novel V and J alleles were inferred using TIgGER. The curated set of alleles from the 106 animals was generated by retaining only DNA genoytpes with sufficient evidence to appear in the AIRR-Seq data for a given animal, as inferred by PIgLET. The resulting data were integrated with existing datasets of IG alleles to form the Macaque Unified Set of Alleles (MUSA). The data is hosted on VDJbase allowing for exploration, summary reports and interactive visualizations comparing genomic and repertoire data. Figure was created in https://BioRender.com

To recover putative alleles, we constructed a “seed” reference set for V, D, and J gene segments across the IGH, IGK, and IGL loci by integrating established sources (KIMDB [6], IMGT [22], and RhGLDB [21]) alongside newly inferred alleles derived from diverse sources (RhGLDB+) and from a *de novo* whole-genome and targeted long-read sequencing assembly of a sire-dam-offspring trios (see Methods section 4.4). Targeted IG long-read genomic data was generated using an optimized protocol adapted from those previously published for the human IG loci [34, 35, 36]. This protocol used a custom target oligo panel designed from known RM IG genomic data to enrich long-read single-molecule real-time (SMRT) Pacific Biosciences sequencing libraries for DNA fragments derived from the IGH, IGK, and IGL loci. Following this approach, sequencing libraries for multiple samples were multiplexed, allowing for high-throughput generation of long-read IG genomic data. High-fidelity (HiFi) long reads were assembled into highly accurate haplotype-specific contigs. The genomic DNA assemblies for each individual animal were then annotated using DIgGER [37] to identify known and undocumented alleles, defining a genomic germline set (GGS) for each subject. The matching data from the AIRR sequence were annotated using these individual GGS’s, applying stringent criteria to identify expressed alleles of high confidence, resulting in an individualized repertoire germline set (RGS) (Methods, Section 4.9). This conservative approach allowed the confident curation of high-fidelity alleles at both the genomic and expressed repertoire levels for each subject.

A summary of the IG alleles detected for each animal at the genomic and expressed repertoire is shown in (Figure 2A). The largest number of alleles were observed for the V genes: for IGHV, a median of 175 alleles (range 89-213) were detected at the genomic level per individual, and 106 (range 57-132) alleles in the expressed repertoire. For the light chains, our approach detected medians of 128 (range 73-154) for IGKV and 104 (range 64-133) for IGLV at the genomic level, and 95 (range 33-116) for IGKV and 77 (range 34-102) for IGLV in the expressed repertoire. The ratio of expressed-to-genomic alleles varied across the three loci: IGKV and IGLV had a median ratio of 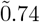, IGHV 0.62, and IGHD 0.63. IGHJ and IGKJ segments showed near-complete overlap (ĩ.0), while IGLJ had a lower ratio (0.6) due to three non-expressed alleles. Across cohort we noted consistent intra-individual overlap of GGS and RGS sequences, supporting the robustness of our approach. We noted a similar degree of overlap between the GGS and RGS when considering the allele sets collectively across all animals for each gene segment type (Figure 2B), in that only a subset of alleles in the cohort-wide GGS were found in the cohort-wide RGS. Together, these findings suggested that the IGH loci harbor a greater number of unexpressed (or lowly expressed) genomic alleles than the IGK/L loci.

**Figure 2:**
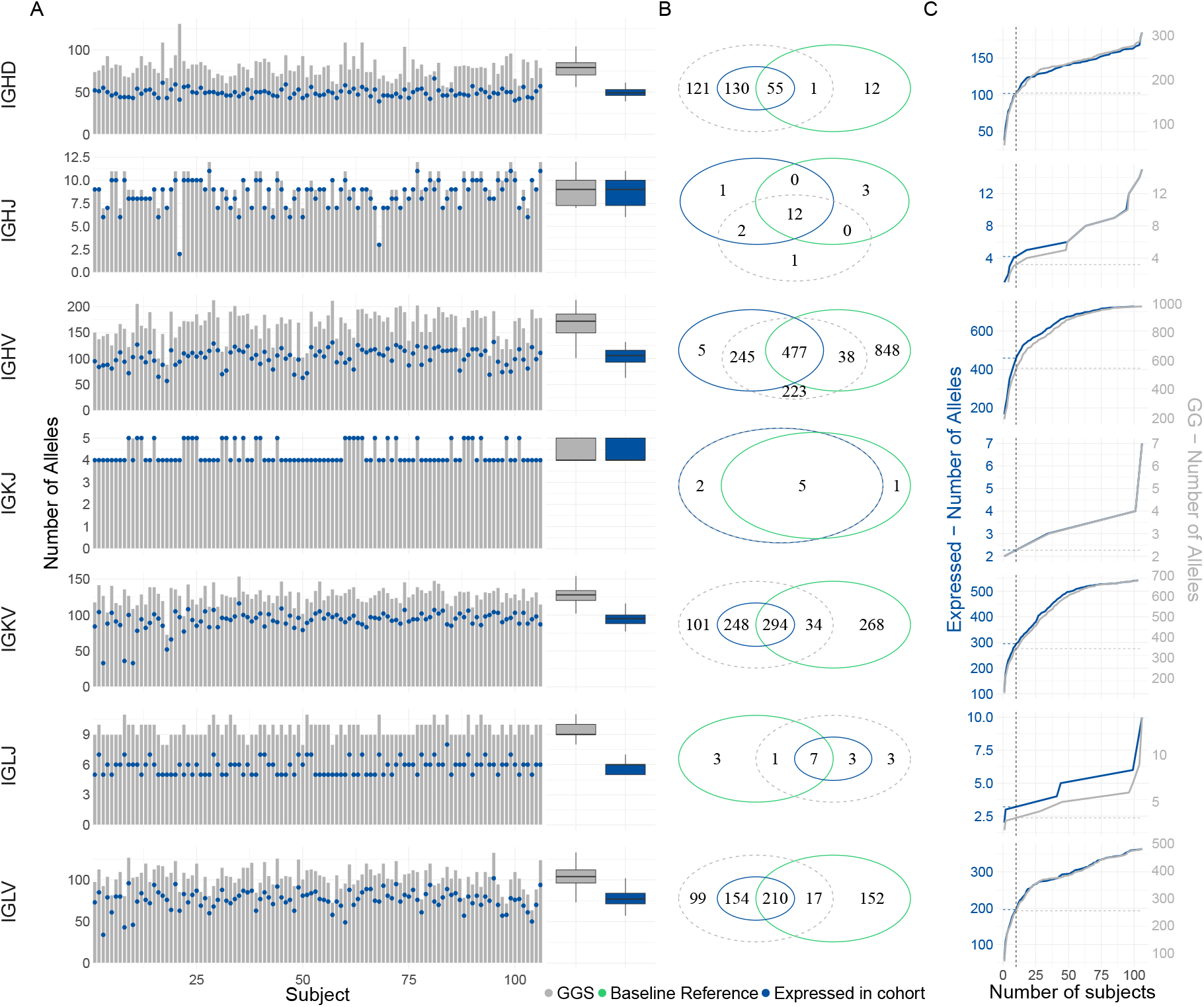
Baseline and De Novo Allele Distribution in Genomic and Repertoire Data of the Current Cohort. (A) The left panel displays allele counts per sample, with the x-axis representing samples and the y-axis showing allele totals. Bars represent allele counts from the Genomic Germline Set (GGS), while blue points indicate Repertoire Germline Set (RGS) alleles identified using the individualized GGS reference. The right panel summarizes the distribution of allele counts across the two reference categories. (B) Venn diagrams showing the relationships between the set of alleles in the Baseline reference (green), the combined alleles from the GGS’s (gray), and the expressed alleles present in RGS annotated using the individualized GGS reference (blue). (C) Cumulative distribution function curves illustrating allele sharing among subjects for each locus. The x-axis indicates the number of subjects sharing a given allele, while the y-axis represents the number of alleles shared by X or fewer subjects. The gray curve represents alleles found in the GGS’s (right axis scale), and the blue curve represents expressed alleles from RGS annotated with the individualized GGS reference (left axis scale).

Notably, there was also only partial overlap in the GGS/RGS sequences with the baseline reference, which comprises previously known alleles, including those from the seed set (see Method Section 4.4) and external sources (Kong et al. [38] and Guo et al. [23]). This highlights previously undocumented alleles found in our cohort, which is investigated in more detail below.

We also observed substantial variability in allelic sets between individuals. For example, among the 727 IGHV alleles expressed in at least one individual, 63% were detected in 10 or fewer animals, reflecting high inter-individual diversity (Figure 2C).

### 2.2 Macaque Unified Set of Alleles (MUSA) Extends the Breadth of All Previously Used Datasets

We created the Macaque Unified Set of Alleles (MUSA) by integrating alleles from the baseline reference described above and previously undocumented alleles newly discovered in this study. This set provides a broader and more comprehensive dataset compared to previously established IG germline databases (Figure 3A). Alleles shared between multiple sources were observed more frequently in both genomic and expressed repertoires than alleles unique to a single source. For instance, within the IGHV locus, alleles found in KIMDB, Trios, RhGLDB+, and IMGT were consistently observed at higher frequencies as expressed in our cohort (70 out of 70) than those exclusive to either source (58 out of 421 for KIMDB, 50 of 73 for the Trios, 8 of 96 for RhGLDB+, and 7 of 137 for IMGT-exclusive alleles). This trend extended to other loci as well; in IGLV and IGKV, all alleles that were shared by more than one source were observed in RGS, as opposed to alleles that were present only in IMGT or only in RhGLDB+ (*e*.*g*., 18 of 163 and 11 of 113 for RhGLDB+ in IGKV and IGLV, respectively). Similarly, for IGHD and IGHJ, alleles exclusive to KIMDB were rarely detected in our cohort (4 out of 9 for IGHD and none for IGHJ), while shared alleles between KIMDB and other sources were more represented within our cohort (32 out of 34 for IGHD and 8 out of 8 for IGHJ, Supplementary Figure 1).

**Figure 3:**
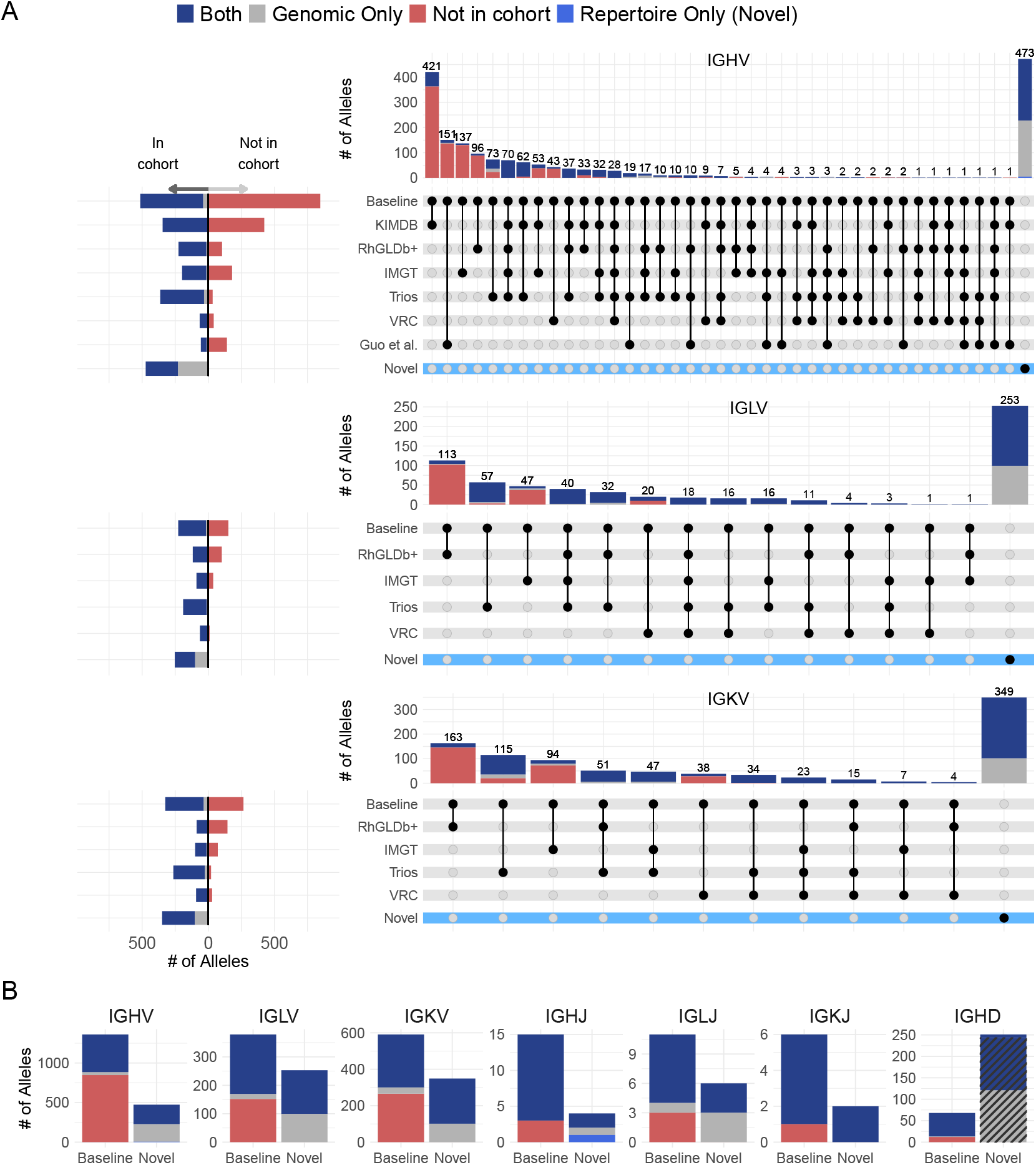
Macaque Unified Set of Alleles (MUSA) Categorized by Source. (A) UpSet plots showing intersections of IGHV, IGLV, and IGKV allele sets derived from various sources. The left-side horizontal bars indicate the number of alleles observed (left) or not observed (right) in the current cohort of 106 animals for each data source. The top vertical bars represent the number of alleles unique to or shared among sources, as indicated by the black dots in each column. (B) Summary of data shown in (A) panels from the UpSet plots shown here (IGHV, IGLV, IGKV) and in Supplementary Figure 2 (IGHJ, IGLJ, IGKJ, and IGHD). Alleles are stratified based on their presence in the baseline reference used for cohort annotation (left) or previously undocumented (right). The bar color scheme for both panels is described in the legend at the top of the figure. The cross lines indicate alleles that were filtered out.

The majority of alleles derived from the Trios dataset were observed in our cohort, both in GGS and RGS, with 50 out of 73 alleles detected for IGHV, 54 out of 57 for IGLV, and 95 out of 115 for IGKV. This likely reflects the similarity in sequencing protocols and the shared origins of the animals used to infer these alleles, leading to a stronger overlap with this larger cohort.

Despite the overlap identified between our cohort and other existing sets, for the majority of loci, newly identified alleles comprise the bulk of alleles curated in the MUSA set (Figure 3B). A significant proportion of identified IGKV (40%) and IGLV (37%) alleles were undocumented, likely reflecting paucity in previously published data for the light chain loci. This was in contrast to IGHV, in which 25% of characterized alleles were previously undocumented. However, we did note a significant increase in the number of IGHD alleles relative to previously published sets. The shorter length of IGHD segments complicates their identification in genomic data, while extensive trimming during V(D)J recombination often obscures their full sequence in repertoire data. Here, we applied a stringent filtration process to overcome these unique challenges. We included only IGHD alleles expressed in the repertoire with a median ratio of observed sequence length to allele length above 0.5 (Supplementary Figure 2). This filtering reduced the total number of IGHD novel alleles from 251 to 8 (Figure 3B IGHD column crossed bar), ensuring that only high-confidence alleles were included.

### 2.3 Unsupervised Clustering of the MUSA Based on Coding Region Sequences Distinguishes Between Expressed and Non-Expressed Clusters

Historical nomenclature systems have relied heavily on the ability to accurately map discovered allelic variants to specific genes within an IG locus. This has traditionally required the assignment of alleles to specific gene labels within a germline set, and for many current AIRR-seq processing pipelines, is deeply integrated into the germline gene assignment process. However, recent observations highlight pitfalls in this approach: 1) in some cases, germline sequences cannot be precisely mapped to a given IG gene; and/or 2) a given germline sequence has been shown to reside at two unique genes within a species. Both of these scenarios can result in erroneous sequence assignments, and ultimately misguided interpretation of gene/allele designations and biological features inferred from the dataset. The IG gene nomenclature for RM is currently in its infancy, and as such, accurate assignment of discovered germline alleles to specific genes is currently not feasible. Thus, to guide the grouping of sequences within the MUSA set, we have applied Allele Similarity Clusters (ASCs) [39], which allows for the grouping of alleles based on sequence similarity cutoffs, rather than genomic position. Critically, this system does not replace, but is used alongside traditional nomenclature systems, allowing for more accurate analysis tasks; additionally, as is the case for our RM set, ASC-based names can serve as interim identifiers in species for which formal nomenclatures are yet to be established.

To construct ASCs for the MUSA germline set, we conducted unsupervised clustering, performed in two steps. First, following Kabat [40], complete linkage clustering was applied with a 75% similarity threshold to define “allele subgroups”. Next, a community detection algorithm grouped alleles into ASCs [39].

The ASCs varied in size, ranging from 1 to 68 alleles (Figure 4A). V segments showed the most diversity, forming clusters of up to 68 alleles in IGHV, 36 in IGKV, and 29 in IGLV. D and J segments were more constrained, with up to 8 alleles in IGHD and 3–4 in the J segments. To further explore the diversity of ASCs and the subgroups they form, we examined the number of ASCs grouped into each subgroup. Subgroups of V segments contained between 1 and 63 ASCs for IGHV, 1 and 61 for IGKV, and 1 and 14 for IGLV. Subgroups of D and J segments were smaller, containing between 1 and 5 ASCs for IGHD and only 1 to 3 ASCs for IGHJ, IGKJ, and IGLJ. Where possible, we maintained correspondence with IMGT subgroup classifications to ensure consistency [22]. To illustrate the distribution of alleles in these subgroups, their sizes were categorized into groups based on allele counts (Figures 4B and 4C). Most ASCs contained only 1–4 alleles, while larger groups with more than 16 alleles occurred predominantly in V segments.

**Figure 4:**
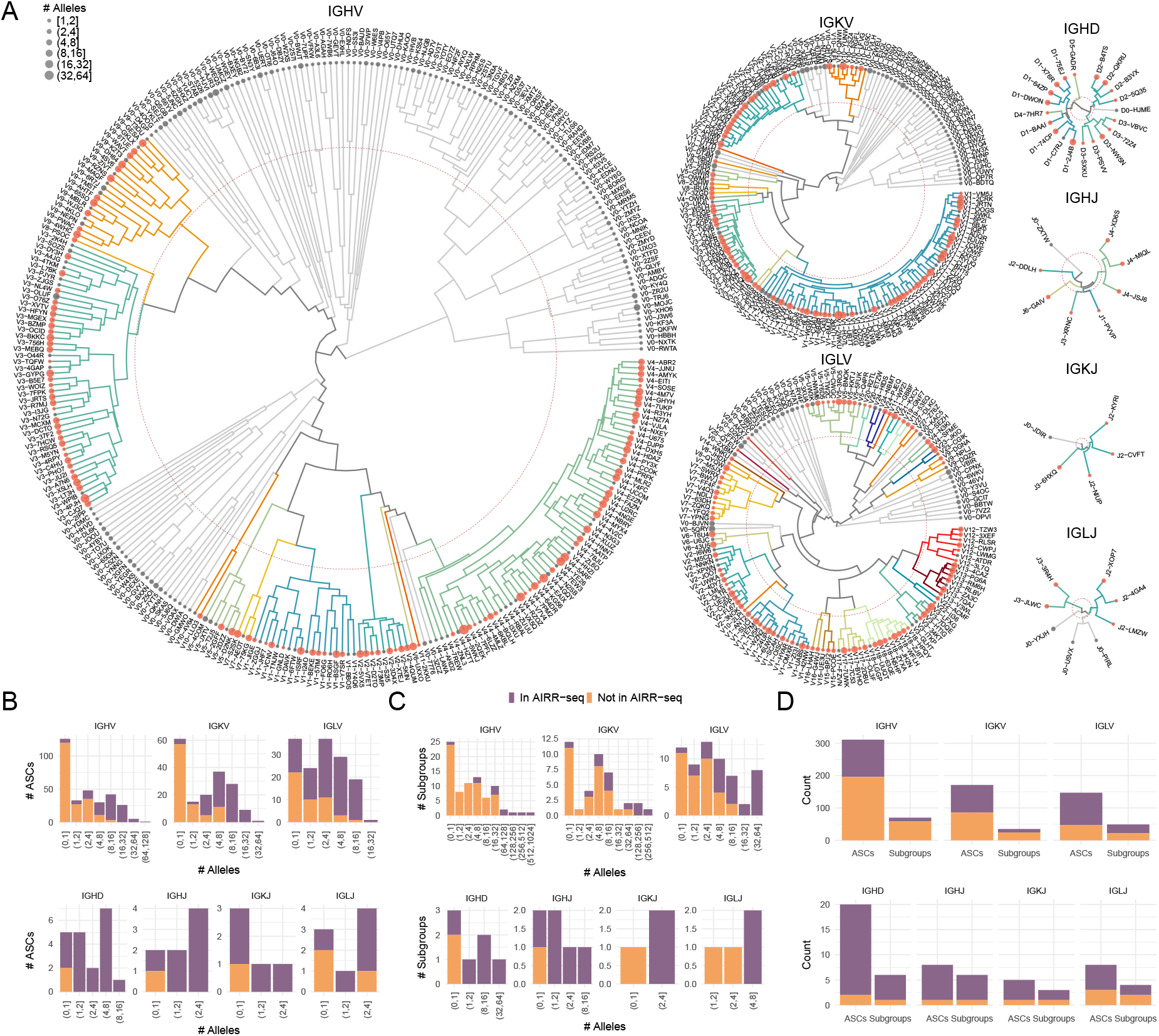
Allele Similarity Clusters (ASCs) within the Macaque Unified Set of Alleles (MUSA). (A) Circular dendrograms illustrate the ASCs comprising the MUSA. Each node represents an ASC, with the dot size indicating the number of alleles in the cluster and the color signifying whether a representative allele of the ASC is present (red) or absent (gray) in at least one repertoire within the cohort. The red dashed circular line marks the 75% similarity threshold used for subgroup definition. Branch colors denote different subgroups, while gray branches represent ASCs not observed in any repertoire and therefore lack subgroup assignments. (B-C) Summary bar plots of the number of alleles per ASC (B) and per subgroup (C) for each immunoglobulin gene type. The x-axis represents the number of alleles grouped into logarithmic bins, indicating the size of each group in terms of allele counts. The y-axis represents the number of ASCs or subgroups corresponding to each allele bin. The color represents whether the ASC or subgroup was seen in the AIRR-seq data (color-coded in the legend). (D) Total ASCs and subgroups in each Ig gene type. The y-axis represents the total count, while the x-axis represents the categories. The color indicates whether the ASC or subgroup was observed in the AIRR-seq data (color-coded in the legend).

A comparison of the fractions of ASCs and subgroups observed in AIRR-seq in our cohort for each gene type revealed notable differences (Figure 4D). The IGHD and IGHJ loci had the highest fractions of ASCs represented in AIRR-seq (approximately 87% and 88%, respectively). In contrast, only 37% of IGHV ASCs and 15% of IGHV subgroups were observed in AIRR-seq, suggesting that a large portion of the IGHV alleles may be non-functional, very lowly expressed, or restricted to a subpopulation that is not present in our cohort. For the IGKJ and IGLJ loci, 67% and 50% of subgroups were represented, respectively, while the IGKV and IGLV loci showed moderate representation, with 31% and 55% of their subgroups observed in repertoire data of current cohort.

To dig deeper into the diversity of the IG genes, we evaluated the number of alleles observed for each ASC in each animal and across the cohort as a whole. Although ASCs are not “genes”, this assessment can be used to approximate gene zygosity states (homozygous vs. heterozygous) within an individual. Zygosity profiles were highly variable throughout the population (Figure 5, and Supplementary Figure 3). For approximately three quarters of the ASCs, we observed 0, 1 or 2 distinct allele sequences in each individual, most likely representing gene deletions, homozygotes/hemizygotes, and heterozygotes, respectively. For the remaining ASCs, we observed animals in which the allele number was *>* 2, indicating that these ASCs represented gene duplication events resulting in gene copy number variants (CNVs). For example, we observed 7 alleles in a single animal for one ASC at the IGKV locus. Relative to IGHV and IGKV, IGLV had fewer ASCs with *>* 3 alleles, indicating that there may be fewer gene duplication expansions in IGL at the population level.

**Figure 5:**
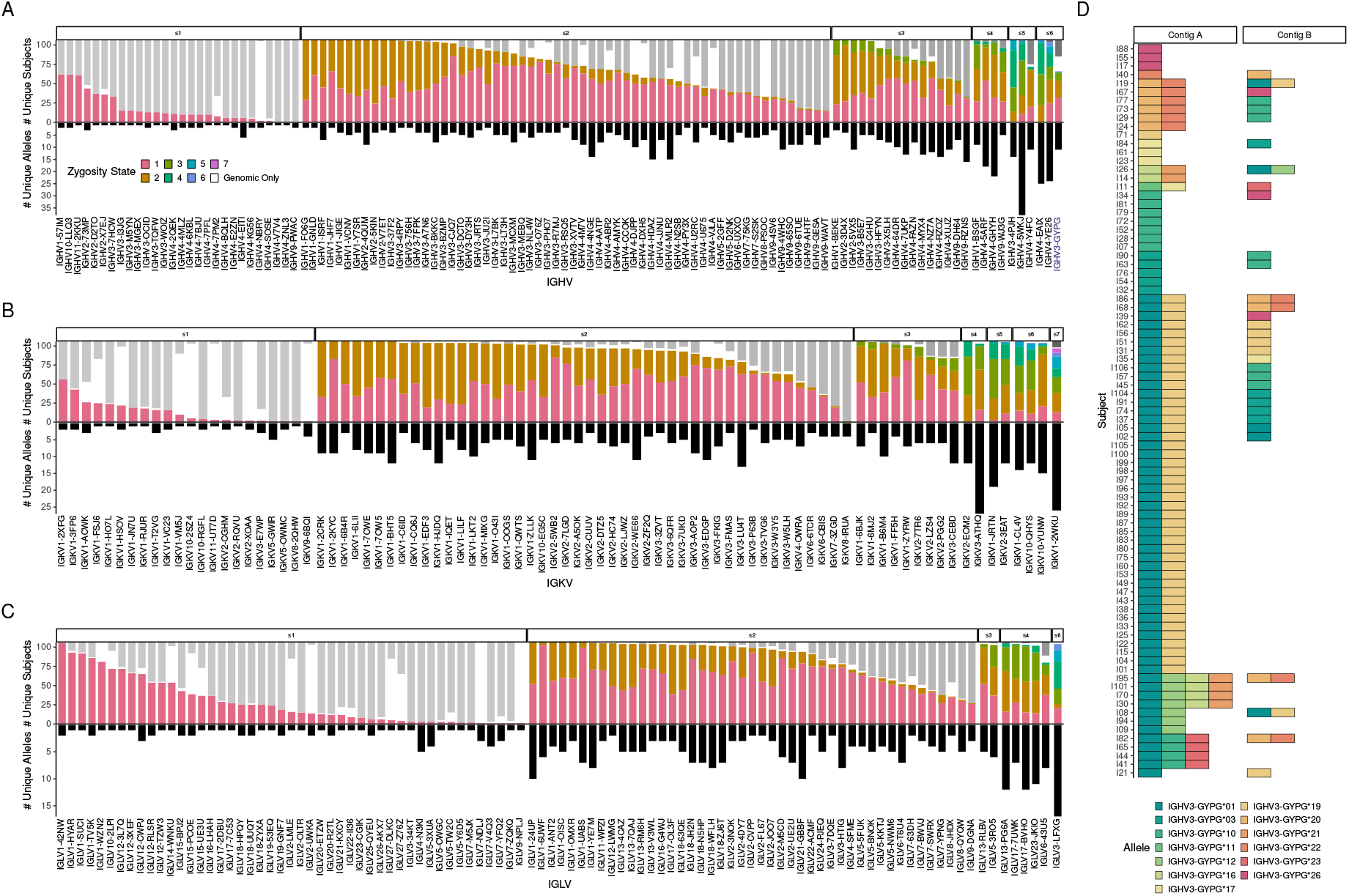
Zygosity Events of Allele Similarity Clusters (ASCs) Across the Population. (A-C) Bar plots illustrating zygosity events for the IG gene types IGHV, IGLV, and IGKV. The x-axis represents different ASCs for each IG gene type, while the top y-axis shows the number of subjects associated with each zygosity event, and the bottom y-axis displays the number of unique alleles detected in AIRR-seq data. The bar colors correspond to zygosity events, as indicated in the legend. (D) A tile plot showing the distribution of alleles within the ASC IGHV3-GYPG (Highlighted in blue in panel A) in genomic contigs across the population. The y-axis represents individual subjects, with each tile indicating a detected allele. The tile colors correspond to specific alleles, as denoted in the legend.

We next examined the genomic assembly data to garner additional support for the presence of CNVs (Figure 5D). Taking, as a case study, ASC IGHV3-GYPG, we noticed that multiple alleles often occurred within the same contig, indicating their presence on the same chromosome and therefore providing evidence of gene duplication. Within each sample, we identified all IGHV3-GYPG alleles found on assembled contigs, noting either one or two contigs per animal (labeled contig A and contig B in Figure 5D). We observed between 1 and 4 alleles per contig, with a range of 1 to 6 total alleles observed per animal. Because assemblies have not been fully-phased into full-length haplotypes, we cannot determine at this point whether when two contigs are observed, they represent alleles residing on different homologous chromosomes. Nonetheless, these results clearly demonstrate the presence of extensive IG gene CNVs.

### 2.4 Analysis of Recombination Signal Sequences of Expressed Alleles Reveals Striking Patterns of Diversity

RSSs are critical genomic elements for the proper functioning of V(D)J recombination and are common motifs for defining “functional” IG genes. However, few studies have had the opportunity to assess the variation in RSSs among IG alleles with evidence of expression in the repertoire within the same animal. Here, we curated RSSs from genomic assembly data for all IG alleles identified in our cohort, including heptamer, nonamer, and spacer elements. We limited our analysis to alleles with high-confidence RSSs, defined as cases where a given RSS was the only non-coding variant for an allele expressed in at least one subject (Figure 6A). We found that RSSs exhibit substantial variability among expressed alleles, gene segment types and loci (Figure 6). In total, we identified 328 RSSs across loci: 78 in IGHV, 12 in IGHJ, 36 in 5’ IGHD, 50 in 3’ IGHD, 71 in IGKV, 7 in IGKJ, 74 in IGLV, and 9 in IGLJ.

**Figure 6:**
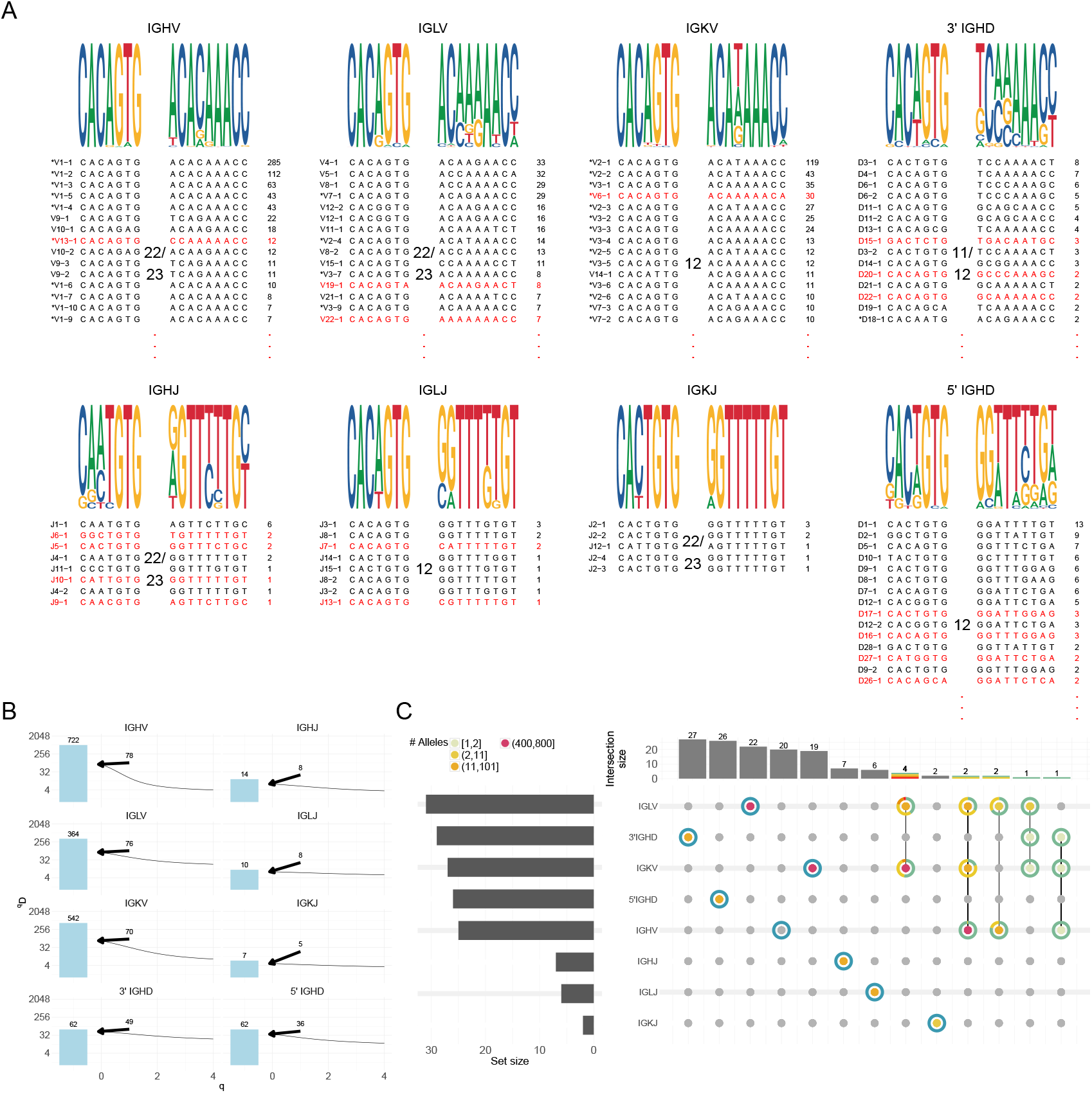
Variability in Recombination Signal Sequence (RSS) Heptamer and Nonamer. (A) Sequence logos (top) represent the variability of RSS heptamers (left) and nonamers (right) for each gene segment, weighted by the number of unique allele-RSS sequence pairs. The degeneracy of each RSS (bottom), measured by the count of unique allele-RSS sequence pairs, is displayed to the right of each heptamer-nonamer sequence. Each RSS is labeled on the left, with an asterisk indicating that the heptamer-nonamer sequence also occurs in other loci. Heptamer-nonamer sequences not observed in humans are highlighted in red. (B) Plots illustrate the number of unique alleles (bar) and the Hill diversity index (line) of the RSS groups for each gene segment. The y-axis uses a logarithmic scale, while the x-axis represents the Hill diversity index parameter *q*. Labels denote RSS richness (unique counts) at *q* = 0. (C) UpSet plots depicting intersections of shared heptamer-nonamer pairs among various loci. The horizontal bars on the left represent the number of unique heptamer-nonamer pairs for each locus. The vertical bars at the top illustrate the count of shared heptamer-nonamer pairs among loci, as indicated by the colored dots within each column. The color of the dots corresponds to the number of alleles carrying the shared heptamer-nonamer pair. Surrounding the colored dots, pie charts provide a visual representation of the fraction of alleles from each locus that contain the shared heptamer-nonamer pair.

In addition to assessing sequencing variation within nonamers and heptamers, we also identified variation in spacer lengths among expressed alleles, including non-canonical lengths. For example, we observed a 22-nucleotide spacer in IGHV associated with four unique heptamer-nonamer pairs, representing 41 coding alleles. Similarly, we observed an 11-nucleotide spacer in the 3’ IGHD, observed for one RSS for a single allele, found in 81 subjects. While most RSS heptamer-nonamer pairs were shared with those in humans, a small subset were unique to RM. For instance, IGHV contains 62 pairs shared with human data from VDJbase, and 16 that were only found in our RM dataset. In IGKV, we identified 51 shared pairs and 19 that were only in RM. Notably, all IGKJ pairs identified in this study were observed in humans.

We found that the RSS spacer exhibited greater variability between loci, relative to intra-locus differences observed for the heptamer and nonamer regions (Figure 6B). Relative to the number of unique IGHV alleles observed, we observed relatively low variability in RSSs. In contrast, the J and D segments exhibited a nearly one-to-one ratio of alleles to unique RSSs. This suggested that RSS specificity is likely critical to guiding the precise recombination and pairing of particular V’s, D’s, and J’s during V(D)J recombination, whereas RSS variation in IGHV may play a less prominent role in dictating variation in V gene selection by recombination machinery.

The locus-specific nature of RSSs was further emphasized by patterns of sharing of heptamer-nonamer pairs across loci (Figure 6C). Most RSSs were exclusive to a single locus. However, IGKV and IGLV displayed an unexpected degree of sharing, with 8 and 9 shared heptamer-nonamer pairs, respectively.

These results highlight the substantial variability and locus specificity of RSSs in the RM population. While the majority of RSS’s heptamer-nonamer pairs in our dataset were shared with those in human, the observed diversity with our dataset, particularly in the J and D segments, underscores the importance of RSS uniqueness in recombination efficiency and the generation of a diverse immune repertoire.

### 2.5 Comparison of RM IG alleles to human IG germlines that encode HIV bNABs

Next, we assessed the sequence similarity of the MUSA to the human V genes that have been described as the unmutated common ancestral (UCA) of known HIV bNAbs [28]; as these genes have direct relevance to the use of RM in preclinical studies of candidate HIV vaccines. For each human IG gene, we report the macaque allele with the highest pair-wise nucleotide sequence identity (Table 1); the top five top RM alleles by sequence identity are provided in Supplementary Table 2. Overall, the mean human-to-rhesus sequence identity was 93.9% for IGHV (n=15), 95.3% for IGKV (n=10), and 94.5% for IGLV (n=11).

**Table 1:**
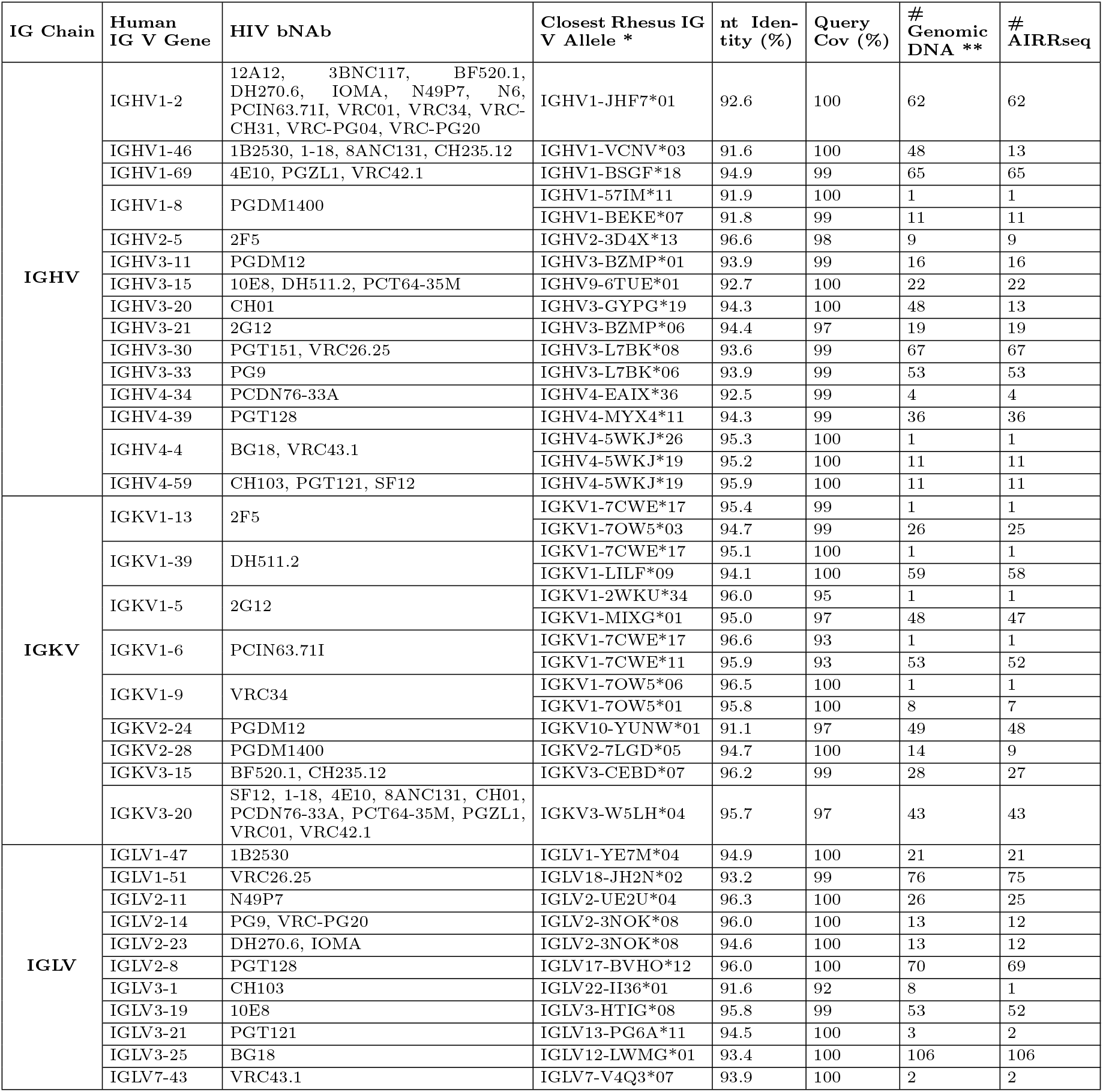
Heavy and light chain V genes in human broadly neutralizing antibodies against HIV and the corresponding closest allele found in Indian origin rhesus macaques in the MUSA dataset. * Only alleles present in both genomic DNA and AIRRSeq data in the cohort of rhesus macaques sequenced for this study are reported in the table. ** In cases where the allele with the highest nt identity was only observed in 1 genome, Table 1 additionally reports representative alleles present at high frequency.

Notably, IGHV1-2, a biomedically important gene in which the UCA is utilized in multiple HIV bNAbs, has generally been considered to lack a close homolog in rhesus. Our results largely confirm this view, as the closest rhesus allele, IGHV1-JHF7*01, had a nucleotide identity of 92.6% with human IGHV1-2, which is only marginally more similar than previous estimates [18]. Similarly, we observed relatively low nucleotide identity for IGHV1-46, in which the closest RM V allele was 91.6%. In contrast, other IGHV genes such as IGHV1-69 (94.9%), IGHV2-5 (96.6%) and IGHV4-59 (95.9%) were found to have relatively high nucleotide identities to the closest human allele. For the light chains, we observed several IG gene segments with $¿ 95%$ identity – IGKV1-13 (95.4%), IGKV1-39 (95.1%), IGKV1-5 (96.0%), IGKV1-6 (96.6%), IGKV1-9 (96.5%), IGKV3-15 (96.2%), amongst others.

This analysis also allowed us to query the prevalence of a given allele within our cohort of 106 RMs. In some instances, such as putative homologs for IGHV1-2 and IGHV1-69, observed occurences were in as many as 62 and 65 animals, respectively. In contrast, for other alleles, such as IGKV1-13, the closest allele, IGKV1-7CWE*17 (96.1% nucleotide similarity), was observed in only one animal. In general, these data allow for detailed insight into the immunogenetics of biomedically important IG alleles in rhesus macaque.

### 2.6 Allele Browser

We have developed an interactive web server, accessible at https://vdjbase.org/referencebook/Rhesus Macaque, that provides comprehensive insights into RM IG data. For each IG chain, the platform offers an Allele Visual Summary, presenting a detailed overview of alleles categorized by gene types, subgroups, and ASCs. Accompanying this summary is a matching Allele Table that includes information on the coding region, such as nucleotide sequences, IgLabel [41], and the other sources where the allele appears, as well as details on non-coding regions, including RSS and leader sequences. Additionally, the web server presents ASCs and the absolute frequencies of alleles within the dataset, along with the chosen allele-specific thresholds. Interactive graphs enable users to explore available group information, thresholds, frequencies, GGS and RGS, and zygosity data by selecting individual data points. This user-friendly interface enhances the analysis of RM IG data, allowing for detailed exploration of allele distributions and relationships.

## 3 Discussion

This study presents the first large-scale resource of IG-encoding variants in RMs, integrating high-quality genomic data with expressed repertoires. From 106 Indian-origin RMs, we identified numerous “novel” alleles across seven IG gene types and loci. Additionally, we curated introns, leader sequences, and RSSs for all identified alleles. These data provide a foundation for developing RM IG germline databases, with broad implications for antibody-mediated immunity research.

The most practical contributions provided by these data stem from the drastic expansion of cataloged germline IG alleles. Germline databases are a mainstay for the investigation of the Ab response via the analysis of AIRR-seq data, facilitating the assignment of sequencing reads to known germline alleles. This is critical for accurately characterizing SHM and delineating Ab lineages and key residues most relevant to the antigen and phenotype under study. Notably, germline databases that better represent standing diversity within a species improve the accuracy of such analyses [42]. Our dataset more than doubles the number of validated RM V, D, and J alleles, and is thus expected to immediately augment both bulk and single-cell AIRR-seq studies. In addition, we have established the use of ASCs for our dataset [39], which allows us to account for uncertainty in gene and allele relationships in the germline assignment process, avoiding error-prone gene assignments. This is particularly important for alleles that are shared among complex gene duplicates, in which false assignments can result in misinterpretation. Given the complexity of the RM IG loci and our incomplete understanding of locus structure and gene homology, ASCs offer a solution for maintaining order and consistency in gene assignments between individual animals, both within and between studies. This will be critical for the optimization and future development of AIRR-seq based genotyping algorithms. Furthermore, relative to other existing germline reference sets, we specifically improved documentation of leader sequences confirming the proper delineation of L-PART1 and L-PART2 by verifying the expressed leader sequence in AIRR-seq. These gene elements are essential for the proper function of IG sequence assembly steps in single-cell AIRR-seq analysis pipelines, and can also be used to review and optimize existing AIRR-seq primer sets and molecular protocols for more representative and efficient PCR amplification.

Beyond allele curation, this study establishes a framework for compiling and sharing RM IG data. All supporting genomic and expressed repertoire evidence is accessible via VDJbase [43], allowing users to explore IG data at both sample and population levels. Identified germline alleles have been annotated with overlaps from external databases (see Supplementary Table 3). Users can download the full dataset for alignment or apply filters to prioritize alleles with demonstrated expression or matched RSS sequences. Some alleles, not observed in our samples, are included and annotated accordingly, with evidential standards varying by source. Recognizing the need for standardized and widely adoptable germline sets, we plan to collaborate with the AIRR Community and the International Union of Immunological Societies to refine these resources and ensure the application of appropriate evidential standards [41, 44]. This will enable the real-time incorporation of newly discovered alleles and regular updates to continually expand and refine these datasets.

With this goal in mind, despite the breadth of our dataset, it is evident that current collections of alleles have not reached saturation, and new genes and alleles will be identified with the inclusion of additional animals. Our future efforts will focus on expanding this resource to include more diverse populations of RMs, as genetic diversity across colonies and geographic origins remains under-explored. Here, we showed that drastic differences between NHP colonies exist, and that broader sampling is critical to more fully characterize the extent of diversity present in captive RMs. This also stresses a newfound need to consider IG genetics as a key biological variable when utilizing and comparing Ab data from monkeys sourced from different colonies. Additionally, future iterations of this resource will focus on integrating functional data alongside curated genes and alleles, particularly pertaining to links between IG variants, Ab structure, repertoire gene usage and function. These links will be key for leveraging knowledge of IG diversity in the RM to inform our understanding of disease, and the development of better therapeutics and vaccines.

A primary rationale for our cataloging of IG alleles at scale was to enhance the use of the macaque model for HIV vaccine studies, in particular those studies testing germline-targeting immunogens. In this regard, the data contained in the MUSA, and in the sequence alignments with human UCAs of HIV bNAbs, allow a more comprehensive understanding of the similarity of RM IG alleles with biomedically relevant human antibodies. Furthermore, accessible genotype information allows for a priori enrollment of macaques in studies in which IG immunogenetics are likely to be a significant factor and provides the basis for incorporation of these immunogenetic data into colony management strategies. The usability of the MUSA dataset in this respect can be further enhanced as: (i) Positional genetic information and functional analyses will develop so that evolutionary relationship between highly similar alleles can be inferred; (ii) Structural information from known bNAbs (e.g. critical residues) will be carefully studied to assign RM alleles as potential bNAb precursors; (iii) As future additions to the MUSA dataset will include animals from additional colonies, it is likely that IG homologs with higher similarity to human IG genes will be characterized.

Finally, our data highlight that challenges persist in our pursuit of better and more comprehensive germline databases. For one, we need to build a system that can reckon with the genomic complexity of the RM IG loci. The collection of alleles we report here are disconnected (intentionally) from the genome, to avoid pitfalls associated with our poor understanding of IG locus genetic structure; namely the incorrect assignment of genes and alleles. Consistent with previous reports [18, 19, 7], our analyses provided unequivocal support for the presence of highly related paralogs with shared alleles, and additional evidence of gene CNV. Specifically, we found that approximately one-quarter of ASCs represented expanded sets of duplicated genes. Without the utilization of comprehensive genomic datasets, such genes will be difficult to categorize and name using traditional nomenclature approaches. To ultimately understand what role these types of genomic variants play in determining characteristics of Abs and circulating repertoires, it will be crucial for gene and allele names to properly represent biological and genetic concepts. Designing a nomenclature system robust enough to accommodate the addition of numerous new alleles curated from various forms of data, particularly while lacking genomic context will be a non-trivial task. This challenge will require a community-wide effort to facilitate efficient communication about curated IG germline variation. In a similar vein, our integration of expressed repertoire and genomic annotations revealed that our understanding of regulatory elements, in particular RSSs, also remains incomplete. Although the field has traditionally relied on the use of “canonical” RSS motifs to define “functionality” from genomic annotations, our analyses clearly showed that noncanonical RSS can be associated with V, D, and J alleles that are, in fact, utilized in the expressed repertoire. This indicated that current definitions of canonical motifs are antiquated and should be revisited and are currently insufficient to classify the functionality of alleles curated solely from genomic data. This will also need to be considered as we build RM IG germline databases for the future.

In conclusion, this study provides a robust and dynamic resource for immunogenetic research in RMs, addressing key limitations in existing IG germline datasets while offering novel insights into IG genetic diversity. As additional data are incorporated, this resource will continue to evolve, ensuring its relevance to advance our understanding of Ab diversity and its implications for immunology and vaccinology.

## 4 Methods

### 4.1 Sample collection

Blood was collected from RMs under ketamine anesthesia via the femoral artery, following Emory National Primate Research Center’s IACUC policy 355 “Blood Collection”. PBMCs were isolated using BD® Vacutainer™ CPT™ Mononuclear Cell Preparation Tubes (Cat no. 362753) per the manufacturer’s instructions. The isolated PBMCs were washed with DPBS (Cat no. 21-031-CM) in 50mL conical tubes and resuspended in ACK Lysis Buffer (Cat no. 118-156-101) for 10 minutes to lyse red blood cells, followed by additional DPBS washes.

PBMCs were aliquoted at 5 million cells per tube and lysed in 350 µL RLT/BME for RNA extraction or flash-frozen in an isopropanol/dry ice bath for later DNA extraction. RNA was extracted using the RNEasy Mini Kit (Cat no. 74106) with DNAse (Cat no. 79256) treatment, following the manufacturer’s protocol for total RNA extraction. RNA concentration was measured using the RNA BR Qubit Assay (Cat no. Q10210), and quality and RINe scores were assessed using the 4200 TapeStation RNA Screentape.

### 4.2 AIRR-seq Library preparation and sequencing

PBMCs were isolated from blood samples using density gradient centrifugation with Ficoll-paque. The procedure for library preparation was based on protocols provided by Dr. Daniel Douek (NIAID/VRC) that have been previously published [45, 46]. The IgM constant region primers were based on the primer sequence as previously described [17]. The extraction of RNA from PBMCs was performed using the RNeasy Micro-DNase Digest protocol (QIAGEN) on QIAcube automation platforms (Valencia, CA) into 350 µL QIAGEN RLT buffer. The reverse transcription step was performed using Clontech SMARTer cDNA template switching.

A total of 1 µL of 12 µM 5’ CDS oligo(dT) primer was combined with 8 µL of RNA for 3 seconds. After incubation at 72°C for 3 minutes and cooling to 4°C for at least 1 minute, 8.5 µL of master mix was added, comprising 3.5 µL of 5x RT Buffer (250 mM Tris-HCl (pH 8.3), 375 mM KCl, 30 mM MgCl_2_), 1 µL Dithiothreitol (DTT, 20 mM), 1 µL dNTP Mix (10 mM), 1 µL RNAse Out (40 U/µL), 1 µL SMARTer II A Oligo (12 µM), and 1 µL Superscript II RT (200 U/µL).

The plates were sealed and spun briefly, followed by incubation at 42°C for 90 minutes and 70°C for 10 minutes.

The first-strand cDNA thus obtained was purified using AMPure XP beads.

To amplify the cDNA, 19.3 µL was added to 30.7 µL of master mix containing 25 µL of 2x KAPA Real-Time Library Amplification Kit (catalog# KK2702), 0.7 µL of 5PIIA (12 µM) forward primer, and 5 µL of IgM primer (2 µM) reverse primer. The mixture was vortexed for 5 seconds, centrifuged at 2000 RCF for 1 minute, and monitored using real-time PCR. After amplification terminated in the exponential phase, AMPure XP beads were used for purification.

The addition of barcodes and adapters for sequencing required two rounds of PCR amplification. In the first step, 2 µL of 1:10 diluted amplified rearranged b cell receptor was combined with 46 µL of master mix comprising 25 µL of KAPA HotStart ReadyMix (catalog# KK2602), 2.5 µL of SYBR Green (1:10K), 18.5 µL of nuclease-free water, 1 µL of P5 Seq BC XX 5PIIA (10 µM), and 1 µL of P7 i7 XX IgM (10 µM). The mixture was vortexed, centrifuged, and subjected to real-time PCR amplification and purification as described previously.

In the second step, 5 µL of the amplified library was combined with 44 µL of master mix comprising 25 µL of KAPA HotStart ReadyMix (catalog# KK2602), 1 µL of P5 Graft P5 seq (10 µM) forward primer, 18 µL of nuclease-free water, and 1 µL of P7 i7 XX IgM (10 µM) reverse primer.

A final round of purification with AMPure XP beads was performed. The quality of libraries was assessed using an Agilent Bioanalyzer. The libraries were pooled and sequenced as 309-base paired-end runs on an Illumina MiSeq.

### 4.3 Genomic library preparation and sequencing

Targeted long-read sequencing of the immunoglobulin (IG) loci from 100+ RMs was performed using Pacific Bio-sciences single-molecule real-time (SMRT) technology. Sequencing libraries were generated by adapting a published protocol for targeted enrichment of human immunoglobulin loci. A custom oligonucleotide probe panel (‘Hyper-Explore’, Roche) was designed to target the immunoglobulin heavy chain (IGH), kappa (IGK), and lambda (IGL) genomic regions, utilizing sequences from the RM genome reference (RheMac10) [47] and alternative haplotype assemblies [19].

High-molecular-weight genomic DNA was extracted from peripheral blood mononuclear cells (PBMCs) using the DNeasy Blood & Tissue Kit (Qiagen). DNA (1–2 µg) was sheared using g-TUBEs (Covaris) and size-selected with a BluePippin system (Sage Science). The size-selected DNA was subjected to end repair and A-tailing following the KAPA HyperPrep library preparation protocol (Roche) and ligated to sample-specific barcodes and universal adapters. PCR amplification was conducted using PrimeSTAR GXL DNA Polymerase (Takara) with 8–9 cycles. Amplicons were further purified and size-selected using 0.7× AMPure PB beads (Pacific Biosciences). Target enrichment was achieved through hybridization with IGH, IGK, and IGL-specific oligonucleotide probes (Roche), followed by recovery with streptavidin-coated magnetic beads (Life Technologies). A secondary PCR amplification was performed with 16–18 cycles using PrimeSTAR GXL Polymerase. SMRTbell libraries were constructed using the SMRTbell Express Template Prep Kit 2.0 (Pacific Biosciences), incorporating DNA damage repair and end repair steps. Following A-tailing and ligation of SMRTbell adapters, libraries were treated with a nuclease cocktail to remove unligated fragments and purified using 0.45× AMPure PB beads. Final libraries were pooled and sequenced on the Sequel IIe platform (2.0 chemistry) with 30-hour movie acquisitions to generate high-fidelity (HiFi) reads, with an average read accuracy of 99.8.

### 4.4 Seed reference set curation

The seed reference set used to annotate genomic sequences was derived from a database curated by a Working Group of the AIRR Community (all authors are members of the Working Group). The database included sequences from KIMDB [6], IMGT [22], and RhGLDB [21]. As part of this activity, the authors of RhGLDB contributed additional sequences inferred from repertoires, including light chain sequences and VH alleles. In this manuscript, we refer to the augmented RhGLDB database as RhGLDB+. In addition to alleles from the Working Group database, functional alleles were incorporated from whole-genome assemblies (WGS) of nine RMs, denoted the Trio set. The seed reference set included 1054 IGHV, 65 IGHD, and 15 IGHJ alleles for the IGH locus; 523 IGKV and 6 IGKJ alleles for the IGK locus; and 342 IGLV and 11 IGLJ alleles for the IGL locus. KIMDB covered the IGH locus only, whereas other sets contained sequences from all three IG loci.

### 4.5 Genomic Sequencing Processing and individualized Reference Set Construction

#### 4.5.1 Genomic Data Assembly and Filtering

Following genomic data acquisition, de novo assembly was performed using hifiasm [48]. The resulting contigs were filtered based on the following criteria: contigs aligning to the RheMac10 IG loci within chromosomal coordinates using minimap2 [49] (IGH: chr7:167,907,503–169,865,360; IGK: chr13:16,797,737–18,140,859; IGL: chr10:29,621,424–30,895,807), contigs showing significant alignment to validated, high-accuracy long-read assemblies from the Emory RM breeding colony, and contigs exhibiting greater than 70% sequence similarity to the seed set using BLAST [50]. Filtered contigs were categorized into IGH, IGK, and IGL loci groups for further annotation.

#### 4.5.2 Contig Annotation and Allele Identification

Annotation of the filtered contigs was performed using DIgGER [37], with the seed set as reference (see Methods, Section 4.4). DIgGER identified de novo alleles and extracted recombination signal sequences (RSS) heptamer and nonamer elements using predefined position-weighted matrices (PWMs) based on known sequence motifs. Spacer sequences, located between the heptamer and nonamer, were extracted directly from the assembled contigs based on DIgGER-reported coordinates. For antisense strand sequences, the reverse complement of the contig was used to identify spacer positions.

#### 4.5.3 Spacer Length Handling

Spacer lengths vary by gene segment. IGHV, IGHJ, IGLV, and IGKJ segments typically use 22–23 bp spacers, IGKV and IGLJ use 11–12 bp spacers, and IGHD uses a 12 bp spacer. To account for non-canonical spacer lengths, DIgGER was run twice. The first run used canonical spacer lengths, while the second run employed expanded spacer ranges of 10–13 bp and 21–24 bp for V and J segments, and 11–13 bp for D segments. Outputs from both runs were combined, with priority given to alleles identified with default spacer lengths in cases of overlap.

#### 4.5.4 Individualized Reference Set Construction

Individualized reference sets were constructed from DIgGER-inferred alleles. Alleles were included if they had more than 10 fully spanning reads at 100% sequence identity. Under relaxed conditions, alleles were also included if they were part of the seed reference set or observed in another subject with more than 10 fully spanning reads at 100% sequence identity. Alleles classified as pseudogenes or those lacking recognizable heptamer or nonamer sequences were excluded to maintain accuracy and reliability.

#### 4.5.5 IGHD Allele-Specific Criteria

Additional criteria were applied to ensure robust IGHD allele identification. IGHD alleles were retained only if they were located adjacent to the common allele similarity cluster IGHV6-UXXO near the D region if two functional alleles were positioned adjacent to each other in the contig, or if they appeared in a sequence of at least three consecutive IGHD alleles, provided at least one allele was part of the seed reference set.

### 4.6 AIRR-seq Data Preprocessing and Quality Control

The AIRR-seq data pre-processing pipeline consisted of two main steps using the pRESTO toolkit [51]. First, paired-end reads were assembled using AssemblePairs with the align function and default parameters. The assembled sequences were then filtered for quality control using FilterSeq with a Phred score threshold of 20.

For each locus in each subject, pre-processed sequence files from technical replicates were merged and aligned against the subject’s individualized genomic reference set using IgBLAST [52]. To ensure unique representation, duplicate sequences were collapsed, retaining only those with a duplicate count of at least 2.

### 4.7 Novel Allele Inference and Alignment

Novel V alleles were inferred using TIgGER [53], enabling the detection of sequence variation up to nucleotide position 318 in the heavy chain and position 330 in the light chain. Similarly, novel J alleles were inferred using TIgGER with modified parameters, detecting sequence variation up to nucleotide position 46 in the heavy chain and position 38 in the light chain.

The sequences were then re-aligned to a germline reference set updated to include any newly inferred alleles. Clone identification was performed using Change-O’s DefinedClone [54], and a single representative sequence with the fewest mutations was selected for each clone to reduce bias during heavy chain individualized repertoire germline set (RGS) inference, commonly known as genotype inference.

### 4.8 Allele threshold optimization

To refine the RGSs inference, allele thresholds were optimized using the cohort’s coupled genomic and repertoire datasets. Repertoire data were initially aligned to the seed set, providing a baseline for comparison with genomic-inferred alleles. This step ensured consistency between the two datasets before threshold adjustments.

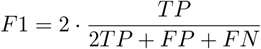

The cohort was divided into training and testing sets, stratified by sequence depth, to improve the robustness of the optimization process. The F1 metric, which balances precision and recall, was used to evaluate thresholds. The F1 score was calculated as:

Definitions for TP, FP, FN, and TN were as follows:

- **True Positive (TP):** repertoire-inferred alleles also present in the genomic inference.
- **False Positive (FP):** repertoire-inferred alleles absent in the genomic inference.
- **False Negative (FN):** genomic-inferred alleles missing in the repertoire-inferred RGS.
- **True Negative (TN):** alleles in the reference not detected in either the genomic or repertoire inference.

A base threshold of 1 × 10^−5^ was applied to all reference alleles, and repertoire allele frequencies were extracted. If an allele was absent, a default frequency of 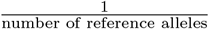was assigned. Z scores were calculated, and thresholds were iteratively adjusted to maximize the F1 score.

Only alleles with Z scores ≥ 0 were considered. When multiple thresholds yielded the same maximum F1 score, the minimal threshold was selected to account for variability in repertoire depth. This process provided a balanced approach to minimizing false positives and negatives in RGS inference.

### 4.9 Subject Filtration and Repetoire Germline Set Inference

To ensure high-quality data for RGS inference and constructing the MUSA, a subject filtration process was applied. Subjects were included only if they had both repertoire and genomic datasets available for all three Ig loci, resulting in an initial set of 132 subjects. Subjects with at least 1,000 annotated reads per chain in the repertoire were retained, reducing the dataset to 125 subjects. Additionally, subjects with low genomic inference coverage or significant discrepancies between genomic and repertoire data were excluded, resulting in a final set of 106 subjects used for RGS inference and MUSA construction.

RGS inference was performed to identify alleles present in each subject’s naive repertoire using stringent filtering criteria. Sequences were required to have no mutations within the V region, a single V allele assignment, and alignment starting at position one of the V germline. For IGHD RGS inference, only sequences with no mutations in the D region and a single assigned D allele were included.

The allele-based method from Piglet [39] was used to determine the presence of specific alleles by comparing allele usage against population-derived thresholds. Briefly, the method compares the allele usage to a population-derived threshold to determine if it should be included in the RGS. The paired cohort allowed for the expansion of the method by including a confidence level metric and fine-tuning the threshold based on a more stable metric (See Methods, Section 4.8).

For each allele inference, a Z score value is calculated. The Z scores assess the significance of an allele’s presence by measuring how its usage deviates from the expected threshold. The higher the score, the more likely that it is truly present in the RGS. The score is calculated as follows:

*N*_*i*_ = Allele count of allele *a*_*i*_ | *a*_*i*_ ∈ *S, P*_*i*_ = Allele frequency threshold, 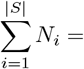 Repertoire depth

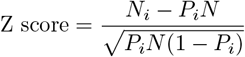

Alleles with Z scores above 0 were included, indicating a high likelihood of true presence in the repertoire.

### 4.10 MUSA Construction and Allele Clustering

The Macaque Unified Set of Alleles (MUSA) was constructed by integrating alleles from the seed set, two external sources (Kong et al. [38] and Guo et al. [23]), and de novo inferred alleles from both genomic and repertoire data. Genomic-derived de novo alleles were included only if they were part of a subject’s “individualized reference,” while repertoire-derived de novo alleles were included only if they met the RGS inference criterion of a Z-score above 0.

To address the unique challenges posed by IGHD segments due to their short length and extensive modifications during V(D)J recombination, a stringent filtering process was applied to de novo inferred IGHD alleles. The ratio of observed sequence length to full allele length in repertoire data was calculated for D annotations matching alleles in the subject’s RGS. Alleles with median ratios below 0.5 were excluded, with this threshold validated against baseline alleles (Supplementary Figure 2).

Novel alleles matching reference alleles from the seed set were replaced with their corresponding references. For alleles detected in both repertoire and genomic data, genomic alleles were prioritized for accuracy.

An Allele Similarity Clustering (ASC) method utilizing a community detection algorithm was applied to group similar alleles. A dissimilarity matrix was constructed using Hamming distance for aligned V allele sequences and Levenshtein distance for unaligned D and J allele sequences.

The dissimilarity matrix was transformed into a similarity matrix by first normalizing the dissimilarity values by dividing each entry by the maximum dissimilarity value. Then, a negative logarithmic transformation was applied to the normalized matrix, scaled by its maximum value:

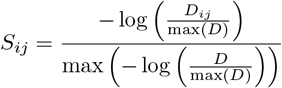

where *D*_*ij*_ represents the dissimilarity between alleles *i* and *j*. Non-finite values resulting from the logarithmic transformation were replaced with zeros. This similarity matrix was then used to construct a weighted graph, where nodes represented individual alleles, and edge weights corresponded to similarity scores between them. Diagonal values were excluded to avoid self-similarity effects.

Combined with the Constant Potts Model (CPM) objective function, the Leiden community detection algorithm was applied to identify ASC clusters [55]. Referred to as Leiden-CPM, this approach addresses the resolution limit inherent in modularity-based clustering, allowing for the detection of smaller, biologically relevant clusters while ensuring cluster connectivity and stability. The Leiden-CPM optimizes community cohesion and performs robustly in complex networks by iteratively refining clusters.

The resolution parameter—a key factor controlling clustering granularity—was systematically optimized to determine the optimal clustering structure. A baseline resolution was first identified by hierarchical clustering, using thresholds of 95% similarity for *V* segments and 90% similarity for *D* and *J* segments. This resolution served as a starting point for further exploration.

A range of resolution parameters around the baseline was surveyed to capture the clustering variability. For each resolution, clustering was performed 100 times with different random seeds to account for stochastic variability in the Leiden algorithm. The clustering configuration that appeared most frequently across the 100 iterations was selected as the representative clustering at that resolution, ensuring stability.

Cluster quality was assessed using the silhouette score, which measures intra-cluster cohesion versus inter-cluster separation. Resolutions producing invalid clustering (e.g., a single cluster) were assigned a silhouette score of −1. The optimal resolution was determined as the one maximizing the silhouette score, yielding the final clustering result.

### 4.11 Incorporating RSS Variation

After defining the MUSA, variations in RSS were analyzed and integrated. Each allele was assigned an RSS-allele pair and categorized as “sure” or “non-sure” based on genomic data. If a single RSS-allele pair was identified in a subject, it was included and categorized as “sure,” regardless of whether the spacer was canonical or non-canonical. If multiple RSS-allele pairs were identified for the same allele in a subject, and all pairs had either canonical spacers or all non-canonical spacers, all pairs were included, but the allele was marked as “non-sure.” If an allele appeared with both canonical and non-canonical spacers within a subject, the pair with the canonical spacer was prioritized.

### 4.12 Alignment of human and rhesus IG genes

The human V gene allele nucleotide sequences were downloaded from OGRDB [44]. These sequences were used as queries against a subset of the MUSA with sample count genomic and sample count AIRRseq *>* 0 (722 IGHV, 542 IGKV and 364 IGLV alleles). Blastn v 2.14.0 [56] was used to find rhesus macaque alleles closest to the human V alleles. The rhesus allele with the highest blastn bitscore among all human allele queries for the given gene was selected for each human gene.

## 5 Funding

The study was supported by the National Institutes of Health [grant number R24AI162317, and NIAID grant number U24AI177622]. Funding for sample collection: Emory National Primate Research Center [P51OD011132]; Next generation sequencing services were provided by the Emory NPRC Genomics Core which is supported in part by NIH P51 OD011132 and NIH S10 OD026799.

## Notes

https://vdjbase.org/

### Competing Interest Statement

The authors have declared no competing interest.

### Summary of Updates

Added one table and edited text

