## supplementary figures for "A Broad Survey and Functional Analysis of Immunoglobulin Loci Variation in Rhesus Macaques"

### Supplementary information for: A Broad Survey and Functional Analysis of Immunoglobulin Loci Variation in Rhesus Macaques

Ayelet Peres[1,2,*], Amit A. Upadhyay[3,*], Vered Klein[1,2,*], Swati Saha[4], Oscar L. Rodriguez[4], Zachary M. Vanwinkle[3], Kirti Karunakaran[3], Amanda Metz[3], William Lauer[4], Mark C. Lin[3], Timothy Melton[3], Lukas Granholm[3], Pazit Polak[1], Samuel M Peterson[5], Eric J Peterson[6], Nagarajan Raju[3], Kaitlyn Shields[4], Steven Schultze[4], Thang Ton[3], Adam Ericsen[3], Stacey A. Lapp[3], Francois Villinger[7], Mats Ohlin[8,9], Christopher A. Cottrell[10,11], Rama R. Amara[3,12], Cynthia A. Derdeyn[13,14], Shane Crotty[11,15,16], William R. Schief[10,11,17,18], Gunilla B. Karlsson Hedestam[19], Melissa L. Smith[4], William Lees[1,4], Corey T. Watson[4,†,§], Gur Yaari[1,2,20,†,§] and Steven E. Bosinger[3,21,†,§]

[1]Faculty of Engineering, Bar Ilan University, Ramat Gan, Israel

[2]Bar Ilan institute of nanotechnology and advanced materials, Bar Ilan university, Ramat Gan, Israel

[3]Emory National Primate Research Center, Atlanta, GA, USA

[4]Department of Biochemistry and Molecular Genetics, University of Louisville, Louisville, KY, USA

[5]Division of Genetics, Oregon National Primate Research Center, Beaverton, OR, USA

[6]Wisconsin National Primate Research Center, Madison, WI, USA

[7]New Iberia Research Center, University of Louisiana at Lafayette, Lafayette, LA, USA

[8]Department of Immunotechnology, Lund University, Lund, Sweden

[9]SciLifeLab, Lund University, Lund, Sweden

[10]Department of Immunology and Microbial Science, The Scripps Research Institute, La Jolla, CA, USA

[11]Consortium for HIV/AIDS Vaccine Development (CHAVD), The Scripps Research Institute, La Jolla, CA 92037, USA

[12]Department of Microbiology and Immunology, Emory School of Medicine, Emory University, Atlanta, GA, 30322, USA

[13]Department of Laboratory Medicine and Pathology, University of Washington, Seattle, Washington, United States of America,

[14]Infectious Diseases and Translational Medicine Unit, Washington National Primate Research Center, University of Washington, Seattle, Washington, United States of America

[15]Center for Vaccine Innovation, La Jolla Institute for Immunology, La Jolla, CA, USA

[16]Department of Medicine, Division of Infectious Diseases and Global Public Health, University of California, San Diego (UCSD), La Jolla, CA 92037, USA

[17]IAVI Neutralizing Antibody Center, The Scripps Research Institute, La Jolla, CA, USA

[18]Moderna, Inc., Cambridge, MA, USA
[19]Department of Microbiology, Tumor and Cell Biology, Karolinska Institutet, Stockholm, Sweden
[20]Department of Pathology, Yale School of Medicine, New Haven, CT, USA
[21]Department of Pathology & Laboratory Medicine, Emory University School of Medicine, Atlanta, GA, USA
[*]These authors contributed equally to this work
[†]These authors jointly supervised this work

March 5, 2025

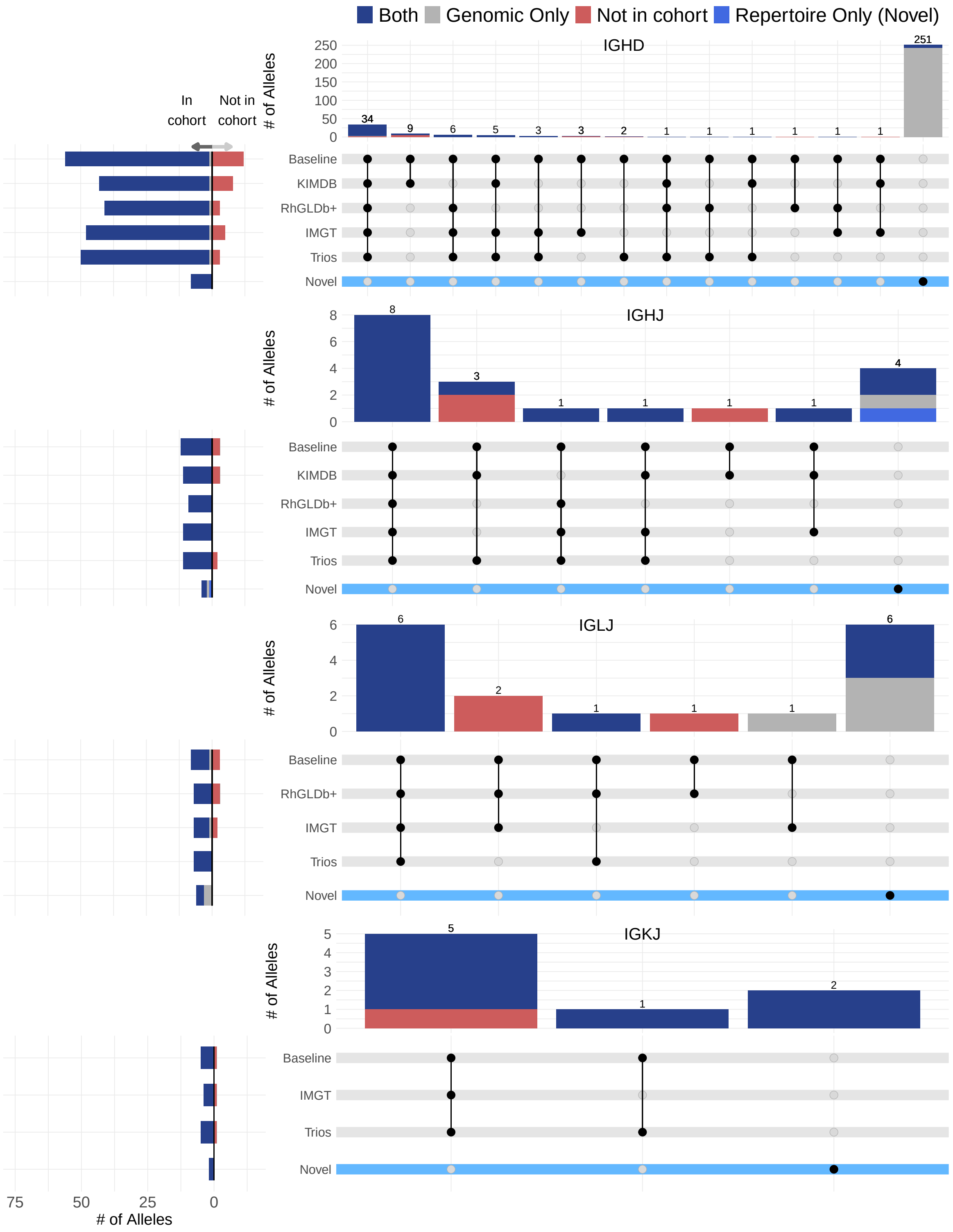


Figure 1: **Macaque Unified Set of Alleles (MUSA) Categorized by Source.** UpSet plots showing intersections of IGHD, IGHJ, IGLJ, and IGKJ allele sets derived from various sources. The left-side horizontal bars indicate the number of alleles observed (left) or not observed (right) in the current cohort for each source. The top vertical bars represent the number of alleles unique to or shared among sources, as indicated by the black dots in each column.

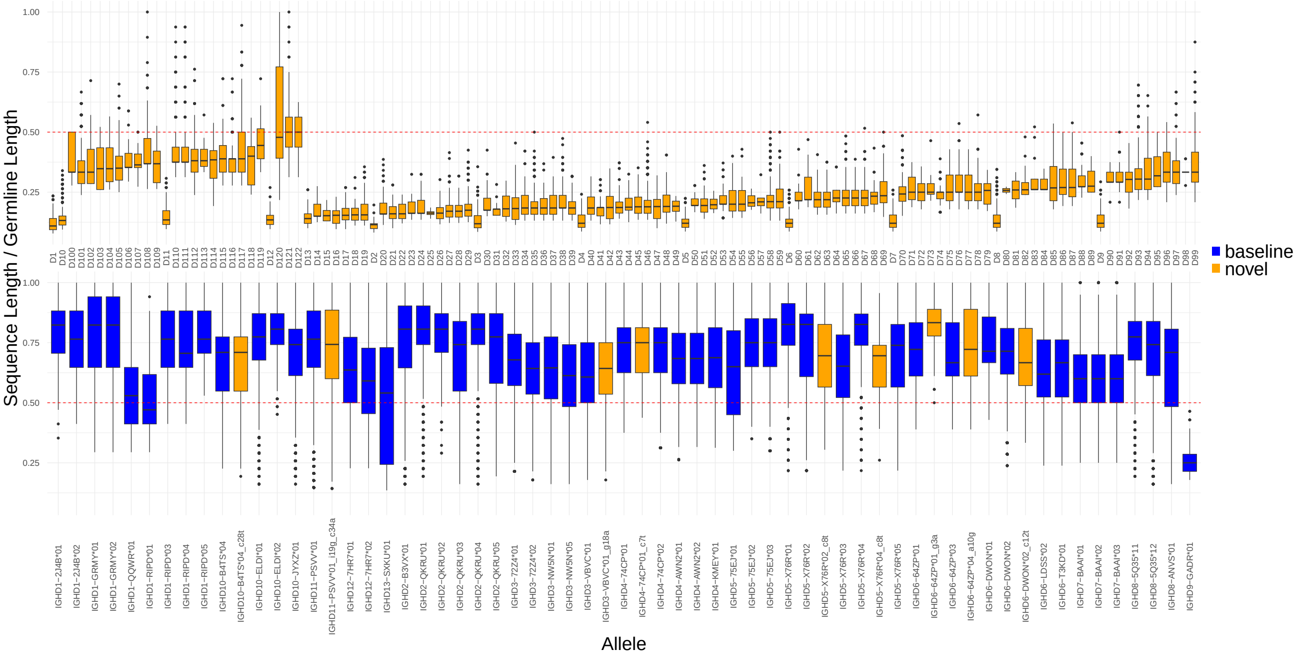


| Label name | Allele | Label name | Allele | Label name | Allele |
|---|---|---|---|---|---|
| D1 | IGHD4−KMEY*01_i0ggaggccgaagcaggcggattacgaggtcaggagatca_t1a_t5c_g8c_g9c_a11g_16acacggtgaaa26 | D42 | IGHD4−KMEY*01_i0tacaggaaatggtca_g2t_g8a_t10a_i11g_17tcatttctttg27 | D83 | IGHD4−KMEY*01_g2t_t5a_g8a_t10a_a11t_g12t_16agcttatt23 |
| D2 | IGHD11−PSVV*01_i0agagattgtaaaatcga_c7a_g11t_a12t_a15t_t17−_a18−_c19−_t22c_c31g_a32c_c33t_35ccttcattaatta47 | D43 | IGHD7−BAAI*01_i0ataatctgcct_g2a_t3c_t5c_t8a_g10t_g11c_c17t_a18t_21ggttttaataac32 | D84 | IGHD4−KMEY*01_g2t_t5a_g8a_t10a_a11t_g12t_16agcttgtt23 |
| D3 | IGHD1−RIPD*01_i0attatcatactcagctgctgctccgatatggactgagta_g1t_t3g_c8t_g10t_g11a_17gcct20 | D44 | IGHD12−7HR7*02_i0ctccttattctctt_g1t_a3g_c4t_g7t_11ctgg14_a18g_a21g_23tgta26 | D85 | IGHD6−64ZP*03_i0c_g1a_g3a_t4c_a5g_a7g_g8c_c12g_c15g_18gtgattgtgc27 |
| D4 | IGHD2−QKRU*04_i0aaggacaaaccacag_c3a_t5g_8ccgc11_t13a_g17−_g18−_c28a_32tgctcctccagagc45 | D45 | IGHD1−2J4B*01_g1t_g2t_t5a_c8t_t9c_g10a_a13t_c14g_t15c_18tcttttttactgcttggaagg37 | D86 | IGHD1−2J4B*02_i0agtag_g1a_g7t_a12c_a13c_a16c_18tcca21 |
| D5 | IGHD2−QKRU*04_i0aaggacgaaccac_c3a_a4g_t5a_7gatcc11_t13c_g15a_t16g_g18c_c28a_32tgctcctccagagc45 | D46 | IGHD12−7HR7*01_t2c_t5−_g6a_g9a_t10a_a15t_a18t_t19g_23tctccacattattggg38 | D87 | IGHD1−RIPD*03_i0gagtgg_g1t_t3g_a4t_a6t_g7c_g11c_a12c_c14a_18gct20 |
| D6 | IGHD2−QKRU*04_i0aaggacgaaccacag_c3a_t5g_8ccgc11_t13a_g17−_g18−_c28a_32tgctcctccagagc45 | D47 | IGHD4−ZLQB*01_i0caattcttgccta_g2t_c7t_c8t_t10a_c13t_16gagatagggtcttactctttca37 | D88 | IGHD4−KMEY*01_i0at_t1a_g2t_a3g_t5a_a11t_c13a_t14g_17atga20 |
| D7 | IGHD2−QKRU*04_i0aaggatgaaccacag_c3a_t5g_8ccgc11_t13a_g17−_g18−_c28a_32tgctcctccagagc45 | D48 | IGHD4−74CP*02_i0tctgaagtctcattat_t1a_g2a_12ttagtgattttttttc26 | D89 | IGHD1−RIPD*04_i0tt_g1t_a4c_a6c_c8g_g11c_a12c_c14g_a16g_18t18 |
| D8 | IGHD5−X76R*01_i0gagcagtgaactgga_t2c_g3c_g4c_t6g_a9t_g12a_t15g_t20a_t21g_24cagccgctgacatcccgagc43 | D49 | IGHD1−GRMY*01_i0cactgtgg_t3g_a6g_g7a_t9a_g10a_a13t_c14g_18gtaaacctcctctcactaatta39 | D90 | IGHD5−75EJ*01_i0gag_a7c_a9g_g10a_c11a_a13g_g15t_g16t_19cag21 |
| D9 | IGHD5−X76R*01_i0gagctgtgaactgga_t2c_g3c_g4c_t6g_a9t_g12a_t15g_t20a_t21g_24cagccgctgacatcccgagc43 | D50 | IGHD12−7HR7*01_i0cag_c4a_6ttttgt11_t19g_g20t_23agaagtggttt33 | D91 | IGHD6−64ZP*01_i0gaggt_a5c_7gaat10_g14t_g15t_18ag19 |
| D10 | IGHD3−72Z4*02_i0gtggcttttttc_g1a_t2a_a6c_t8c_a9g_g11t_g17a_t19−_t22a_29gggcatgtgagcaga43 | D51 | IGHD3−VBVC*01_i0gtgaga_g1c_t5a_7agga10_t13g_g15c_t16a_a17t_28att30 | D92 | IGHD4−AWN2*02_i0accacagtg_t1g_g2t_a3g_t5c_g8c_g9a_t17a_20caggc24 |
| D11 | IGHD3−72Z4*01_i0gtggcttttttcaaattcccgtt_t4g_c7g_t10a_g11t_g15c_t19g_t20c_t23g_a24t_c25g_c27g_29ag30 | D52 | IGHD4−KMEY*01_i0tacaggaaatggtca_g2t_g8a_t10a_i11g_17tcatttttttg27 | D93 | IGHD1−2J4B*02_i0gct_g2c_a4c_g7c_c8a_a12g_a13g_a16g_18tct20 |
| D12 | IGHD3−72Z4*02_i0gtggcttttgc_g1a_t2a_a6c_t8c_a9c_g11t_g17a_t19−_t22a_29gggcatgtgagcag42 | D53 | IGHD1−RIPD*04_i0actttattatttacagg_g1c_g2t_t3a_a4t_a7t_g10t_a12t_c14t | D94 | IGHD7−BAAI*01_2aagc5_g11t_14gacg17_20aatt23 |
| D13 | IGHD2−QKRU*04_i0aaggacgaaccacag_c3a_t5g_8ccgc11_t13a_g17−_g18−_c28a_32tgctcc37 | D54 | IGHD5−75EJ*02_i0atgtaaaattctta_g3t_g4a_a5t_a9t_g10a_c11t_c14t_20ga21 | D95 | IGHD6−UTCO*01_i0gct_g2c_g3t_a5t_a7t_g8t_a10c_g11a_a14c_16ccct19 |
| D14 | IGHD1−RIPD*01_i0gacgcaccca_a4t_a6g_a7c_g11a_a13c_g15c_18tgtgactattaca30 | D55 | IGHD6−DWON*01_i0gcaccaga_g1c_g2c_g3c_a5g_a7c_g8c_g11a_t19c_22ggttcc27 | D96 | IGHD1−2J4B*01_i0cttgc_g1t_g2a_t9a_a13g_c14a_t15c_18gc19 |
| D15 | IGHD6−64ZP*01_i0gcttcacattcaagctgtctccagca_t4a_t6g_c9−_g15c_19tct21 | D56 | IGHD1−RIPD*01_i0tgtagtagtga_a4c_t5a_a7t_g11a_a12g_a13c_18agaagt23 | D97 | IGHD1−GRMY*01_i0gtgg_g1c_a2c_t5g_c8g_g10c_g11t_a12t_c14g_a16t |
| D16 | IGHD2−QKRU*04_i0aaggacgaaccacag_c3a_t5g_8ccgc11_t13a_g17−_g18−_c28a_32tgctcctc39 | D57 | IGHD5−75EJ*03_i0tctcca_g1c_a7c_a9t_c11a_t12a_c14t_t18g_20gagaag25 | D98 | IGHD1−RIPD*04_g1t_a4c_a6c_c8g_g11c_a12c_c14g_a16g_18t18 |
| D17 | IGHD6−64ZP*03_i0ttgctgaggctggaatgcagtggtatgatcata_g2c_g3t_t4c_t6c_a13c_g14t_t16a_19c19 | D58 | IGHD5−X76R*04_i0ccaggt_g1a_t6g_c8a_a16g_c17g_a18g_t20c_22gttcctcttcg32 | D99 | IGHD12−7HR7*02_g1c_t2a_a3g_c4t_g9c_a12g_t16c_a18t_t19a_22aga24 |
| D18 | IGHD6−64ZP*01_i0cttcatggatgtgg_g1a_g2c_t4a_g8t_c9t_a10t_g14t_g15c_19agatttagaaagt31 | D59 | IGHD6−64ZP*03_i0a_g1a_g2t_t4c_a5c_t6a_c9a_t16g_a17g_19cagttgggatccagtgtga37 | D100 | IGHD4−KMEY*01_g2c_a3t_a6t_g9t_a11t_14aggtg18 |
| D19 | IGHD9−GADR*01_i0tccctgaactttcattt_a3c_a4c_g6c_c8g_a10t_a12g_g16c_g20c_t21a_a23c_c24t_g27c_t28a | D60 | IGHD9−GADR*01_t5g_c8a | D101 | IGHD6−64ZP*01_g1c_g3t_a7t_g8c_a10c_g11c_g14t_17ttgct21 |
| D20 | IGHD6−T3KD*01_i0ctggtaccaaaac_g1a_g3a_t4g_g8c_c9a_g11a_a13c_g14a_c15a_22agaacagagc31 | D61 | IGHD1−GRMY*01_i0gta_g1a_a6t_c8g_t9a_a12c_c14t_17aatgcataggtgg29 | D102 | IGHD7−BAAI*01_g2−_t3−_a6t_g7a_a16c_c17g_21taa23 |
| D21 | IGHD3−72Z4*02_i0tttctataatcaggcttagatctta_t2g_a3g_13−−−−16_a21t_t22g_a24c_c25t_29a29 | D62 | IGHD12−7HR7*01_i0caat_a3t_g11−_a12−_a15t_t16a_a18g_t19g_a21c_23tcttttat30 | D103 | IGHD1−2J4B*01_i0aa_g1a_t3c_t5g_a6t_c8g_g10a_g11a_a12g_t15g_18aatt21 |
| D22 | IGHD4−KMEY*01_i0atgaagacgctgtggtgtgtctg_a3t_c4g_a6c_g9t_t10a_g12c_t14c_17agggatgaaga27 | D63 | IGHD12−7HR7*02_g1c_t2a_a3g_c4t_g9c_a12g_t16c_a18t_t19a_22agaaaatatgt32 | D104 | IGHD6−UTCO*01_i0cagacga_g1t_a5c_c9t_a10g_c12g_15gc16 |
| D23 | IGHD10−B4TS*01_i0aaaaagt_6caac9_11−caaca16_c18a_t20c_c28t | D64 | IGHD4−74CP*02_i0atct_a4t_c7g_15ttggtattttccc27 | D105 | IGHD6−LDSS*01_2−−−−5_a7t_g8t_g17t_t19c_22ctg24 |
| D24 | IGHD12−7HR7*01_t2c_t5a_g7a_g9a_g11−_a15t_a18t_t19g_23tctccacattattggg38 | D65 | IGHD4−74CP*02_i0cccggatgacacac_t1a_a3c_t5a_a11g_c13t_16tg17 | D106 | IGHD6−UTCO*01_g1a_g3a_t4g_c9g_a10c_g11t_14cttg17 |
| D25 | IGHD4−KMEY*01_i0tattcctttggacattttt_g2t_a3c_a6t_g8t_g12t_c13t_t14a_17gatattta24 | D66 | IGHD5−75EJ*01_i0tattttatca_g3a_a5t_t6a_a7c_a9t_c11t_t12g_g15t_20tg21 | D107 | IGHD4−KMEY*01_i0t_t1a_t5g_g9−_t10−_g12t_16gtggtagc23 |
| D26 | IGHD5−75EJ*02_i0a_g1a_g3t_g4t_a7t_10aagt13_t17a_a19g_21attctttttactgcttggaagg42 | D67 | IGHD7−BAAI*01_i0gc_g2t_a4c_6tcaa9_g11a_c14a_20tattaaaagc29 | D108 | IGHD1−2J4B*01_t3g_a4g_a6g_11aggg14_17gtt19 |
| D27 | IGHD3−NW5N*01_i0ctgac_g1c_a9g_g15−_g18c_21aaat24_t26g_a27t_31tgcaatc37 | D68 | IGHD1−RIPD*01_g2t_t3g_c8g_g11t_a13c_g15a_17atcttagaatttac30 | D109 | IGHD1−RIPD*01_t3−_t5g_a6g_a7t_g11t_a12t_a13c_17tgta20 |
| D28 | IGHD5−X76R*02_i0cagggcagtgtactagcaca_t6g_a7c_t11a_a16c_a18c_a22−_c23− | D69 | IGHD7−BAAI*03_i0ccatgttattaa_t5a_g7t_t8g_g11a_21aa22 | D110 | IGHD4−74CP*02_t1g_a3g_t5g_a6c_t10a_a11c_a12c_a15g |
| D29 | IGHD1−RIPD*01_i0gacgcaccca_a4t_a6g_a7t_g11a_a13c_g15c_18tgtgactattaca30 | D70 | IGHD4−74CP*02_i0ctgtgcaaatt_t1g_g2t_a3c_a11c_15ttcctcac22 | D111 | IGHD4−74CP*02_t1g_a3g_t5g_a6c_t10a_a11c_a12c_t14c_a15g |
| D30 | IGHD7−BAAI*01_i0ggcaaggttcagaat_g2t_t8g_g9a_g10t_c14a_t15c_a16t_19gttggat25 | D71 | IGHD1−GRMY*02_i0cctgcagtttactg_8taatg12_18t18 | D112 | IGHD1−2J4B*01_g1a_g2c_t3a_t5g_c14t_t15a_a16t_18acac21 |
| D31 | IGHD5−75EJ*01_i0acagtcttgttcct_t2c_g3c_g4t_a7t_t8g_c11a_a13c_g15t_20tggact25 | D72 | IGHD4−KMEY*01_i0gtatttttc_g2c_a6t_g8t_g9t_g12c_c13t_15ttcct19 | D113 | IGHD4−KMEY*01_i0ccttac_g2c_a3t_g12−_t14c_c16a |
| D32 | IGHD2−B3VX*01_i0cctgctgct_g2t_a4c_a6c_c7a_g12−_t13a_18tgact22_32ttgccttg39 | D73 | IGHD4−KMEY*01_i0tcttta_g2t_a3t_g9a_a11c_g12t_15catctcca22 | D114 | IGHD1−2J4B*02_i0acaataa_a13t_c14a_16gctc19 |
| D33 | IGHD3−VBVC*01_i0gcaatattatccttcaa_g1a_a3t_t8−_t10g_g12t_14caat17_c28t | D74 | IGHD4−KMEY*01_i0ttcttttttttctgatgcctt_8tact11_c13g | D115 | IGHD1−GRMY*01_a2−_t3−_c8g_t9a_a12t_c14t_18agg20 |
| D34 | IGHD1−2J4B*02_i0gtgaaagtgag_g1c_g2t_t3g_a4t_c8a_16tccctctcactt27 | D75 | IGHD7−BAAI*03_i0gtgtca_t3g_a6t_g7c_g9a_a12g_a16c_c17t_20gtt22 | D116 | IGHD1−GRMY*01_a2−_t3−_c8g_t9a_g10t_a12t_c14t_18agg20 |
| D35 | IGHD1−GRMY*01_i0atctccatgtaa_g7a_10tact13_a15c_a16t_18cttgccttc26 | D76 | IGHD9−GADR*01_g11a_a19g | D117 | IGHD4−74CP*02_2tgtggg7_a11c_16tat18 |
| D36 | IGHD1−GRMY*01_i0atctccatgtaa_g7a_10tact13_a15c_a16t_18cttgcctttt26 | D77 | IGHD9−GADR*01_t28c | D118 | IGHD12−7HR7*01_g1a_a3t_g6−_g9t_g11t_t16c_a18c_21tttgta26 |
| D37 | IGHD1−GRMY*01_i0atctccatgtaa_g7a_10tact13_a15c_a16t_18cttgctttc26 | D78 | IGHD9−GADR*01_t5g | D119 | IGHD4−74CP*02_i0cagtat_g2a_t10g_13−−−−16 |
| D38 | IGHD4−KMEY*01_i0cgggtaccggtacttact_a3t_a6−_g8t_c13a_17ttttt21 | D79 | IGHD4−KMEY*01_i0tactctg_g2c_a3t_t5c_g8t_g9t_a11g_c13g_17acaattagaaaa28 | D120 | IGHD5−X76R*02_g4a |
| D39 | IGHD4−KMEY*01_i0tgggtaccggtacttact_a3t_a6−_g8t_c13a_17ttttt21 | D80 | IGHD4−74CP*01_i0aaataagcaacatacag_t1a_a3t_c4t_c7t_g9t_c13a_c16t | D121 | IGHD4−KMEY*01_t1a_a3t_c4a_a6t_t7a_t14a_a15g_c16g |
| D40 | IGHD5−X76R*03_i0gattgtatgatag_g1t_g3t_a7t_g10t_t15g_a16c_c17g_t20g_t21g_a22c_24ac25 | D81 | IGHD4−74CP*02_i0gtattatgca_g2c_c7t_g9t_t10a_a12t_15taa17 | D122 | IGHD4−KMEY*01_t1a_g2c_a3t_c4g_a6t_t7a_t14a_a15g_c16g |
| D41 | IGHD1−GRMY*02_i0atgcc_g1t_t3c_a6g_g11t_a13t_c14t_a16c_18aggagaccaaatctctgataa38 | D82 | IGHD1−RIPD*01_i0t_g2t_a4t_c8t_10tttg13_17gctttggg24 | | |

Figure 2: **Analysis of IGHD Alleles.** The boxplots represent IGHD alleles, with the x-axis indicating individual alleles and the y-axis displaying the ratio of sequence length to germline length. Each point within the boxplots corresponds to a sequence annotated with the respective D allele. The top row highlights novel alleles with a median sequence length-to-germline length ratio of less than 0.5. The second row includes seed alleles and novel alleles with a median ratio greater than 0.5. Below the boxplots, a table provides the full names of the novel alleles.

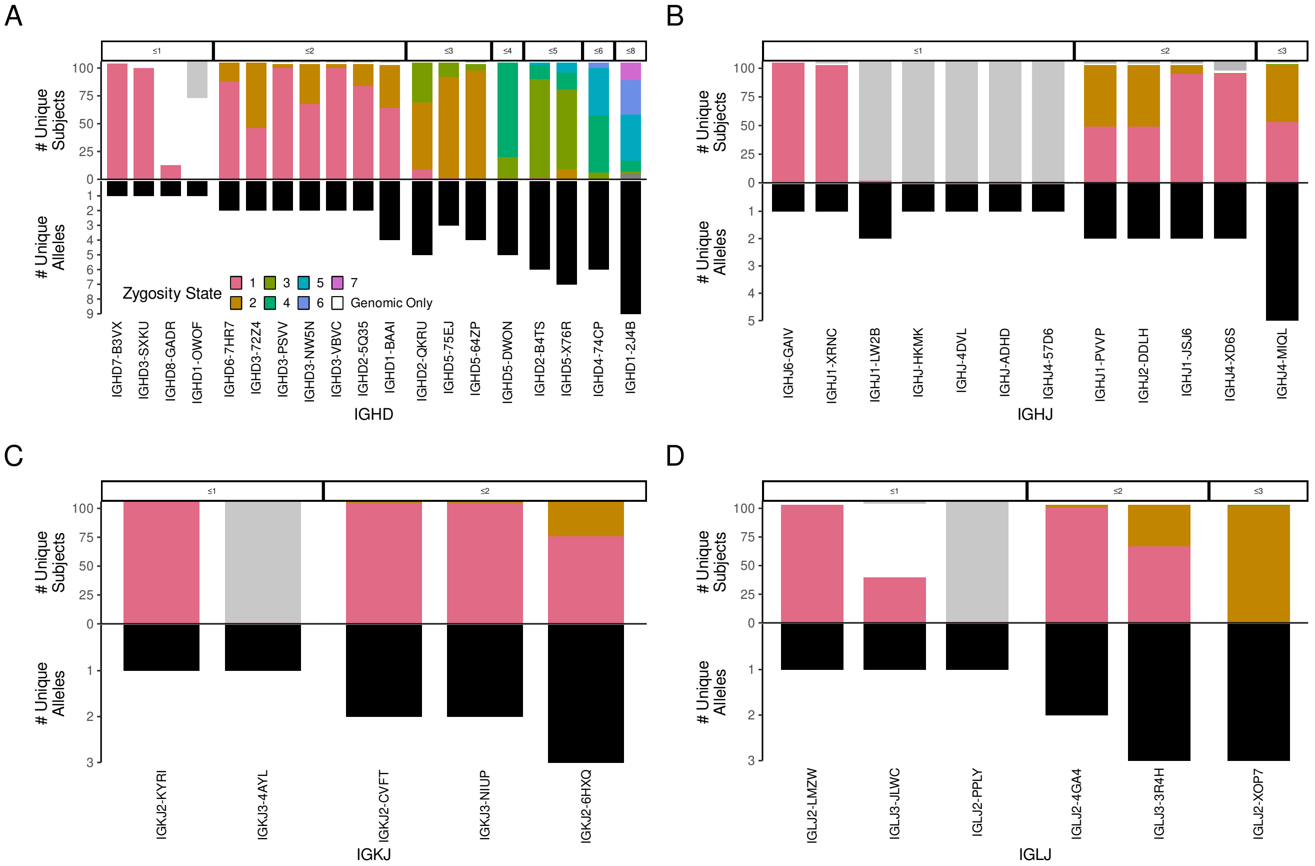


Figure 3: **Zygosity Events of Allele Similarity Clusters (ASCs) Across the Population.** (A-D) Bar plots illustrating zygosity events for the IG gene types IGHD, IGHJ, IGKJ, and IGLJ. The x-axis represents different ASCs for each IG gene type, while the top y-axis shows the number of subjects associated with each zygosity event, and the bottom y-axis displays the number of unique alleles detected in AIRR-seq data. The bar colors correspond to zygosity events, as indicated in the legend.
